# Elucidating cancer cachexia-mediated aberrant cardiac wasting signaling in human iPSC-derived cardiac muscle

**DOI:** 10.1101/2025.08.18.670675

**Authors:** Nora Hosny, Houda Cohen, Mohamed Waleed Elafany, Jeff Schreifels, John Bauer, Rishi Gulati, Brian R. Thompson, Julia C. Liu, Joseph M. Metzger

## Abstract

Cancer cachexia is a highly debilitating clinical syndrome of involuntary body mass loss featuring profound muscle wasting leading to high mortality. Notably, cardiac wasting is prominent in cancer patients and cancer survivors. Cachexia studies present significant challenges due to the absence of human models and mainly short-term animal studies. To address this translational gap, we have developed a robust human-based cachexia experimental approach characterized by marked cardiac muscle wasting and contractile dysfunction, with increased expression of protein degradation markers. Using human iPSC-derived cardiac muscle, we investigated morphological, functional, and metabolic alterations in the key stages of cachexia and in the post-cachexia phase. C26 and HCT116 tumor cell lines were used to induce cachexia by two methods, pulse addition of cancer cell conditioned media or in transwell-adapted co-culture. Cachectic cardiac myocytes exhibited reduced contraction amplitude, prolonged relaxation time, and increased oxygen consumption rate (OCR), as assessed by video-based and Seahorse analyses. Mechanistic investigations centered on the Atrogin-1/Calcineurin A/NFAT axis revealed this signaling pathway as a central driver of cachexia-induced cardiac atrophy. Cachectic cardiac myocytes exhibited significant upregulation of Atrogin-1, leading to a marked decrease in Calcineurin A protein levels. This, in turn, impaired nuclear translocation of NFAT, thereby suppressing its transcriptional activity and downstream cell growth signaling. These molecular changes were accompanied by increased autophagic flux, as indicated by elevated LC3BII/LC3BI ratios. Furthermore, withdrawal of cachexia-inducing stimuli followed by regular media changes for one week led to normalization of Atrogin-1 and autophagy markers; however, functional impairments and metabolic dysregulation persisted, highlighting delayed recovery. Our new findings establish the Atrogin-1/Calcineurin A/NFAT axis as a key regulatory mechanism in cardiac muscle wasting and suggest this aberrant signaling axis may serve as a targetable mechanism for treatment of cachexia-induced cardiac dysfunction.

**Graphical Abstract:** 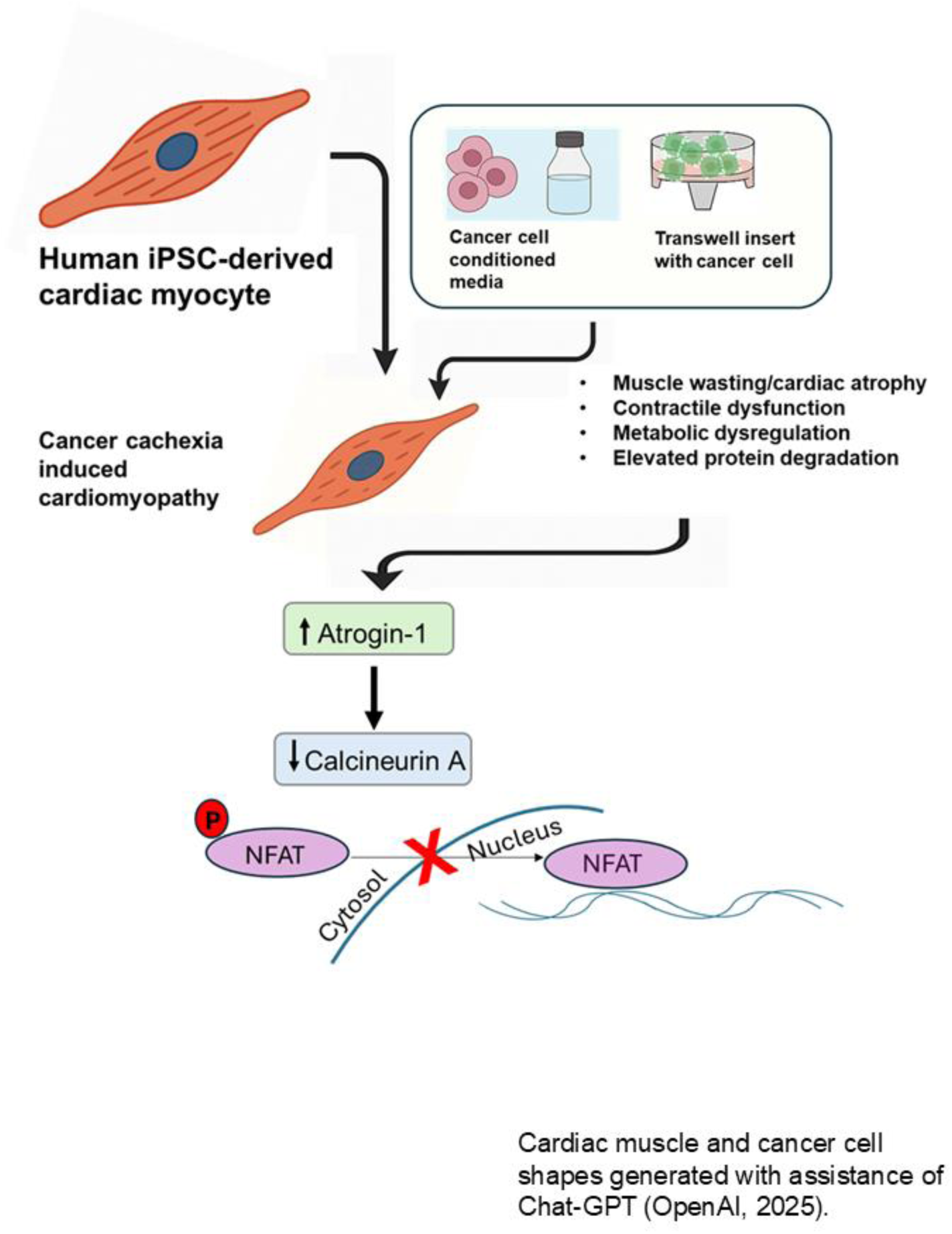

## Introduction

Cardiovascular disorders and cancer, the top two leading causes of death in the United States, present a dual threat to global health affecting tens of millions of patients (1–3). Cardio-oncology represents a new frontier that has emerged as a result of the convergence of these two significant health crises (4). Cachexia is a severe condition, characterized by involuntary skeletal muscle wasting, affecting approximately 80% of cancer patients. Cancer cachexia is directly responsible for the deaths of about 30% cancer patients, often because of heart or respiratory failure (5, 6). With the increasing number of cancer patients worldwide, the burden of cardiac complications in cancer cachexia, namely cardiac atrophy and functional insufficiency, is increasing and has become a significant health issue (3–8).

Chronic cardiac disease sequelae have become increasingly clinically relevant because of improved prognosis of patients with cancer (4, 8). In the United States, there are nearly 20,000,000 cancer survivors, representing 6% of the population. Cardiovascular disease, surpassing cancer, is the leading cause of death among this population (4, 8). Owing to the ever-expanding oncological therapies being introduced into clinical practice, cardio-oncology has traditionally centered on drug toxicities (4); however, this approach has to be expanded to encompass the cardiac insult caused directly by cancer itself, uncoupled from drug toxicities (8). This expanded understanding not only deepens insights into the underlying pathophysiology of the disease but also positions the field for oncological therapies with minimal or no cardiotoxic effects.

Recent studies have reported that the poor cardiac status of patients may intensify the process of cancer and cancer cachexia progression (9–11). Thus, a bi-directional relationship between cancer and the cardiac system exists. In short, cancer cachexia causes cardiomyopathy, and cardiomyopathy can exacerbate cancer progression (12, 13). This vicious cycle culminates with reduced cardiac output, heart failure, diminished quality of life and increased mortality (8). Addressing cancer cachexia-induced cardiomyopathy is crucial to improving outcomes and longevity in both active cancer patients and in cancer survivors.

Although much has been learned about cancer cachexia progression, the processes leading to cardiac atrophy induced by cancer are understudied. This gap in knowledge presents an obstacle to the development of therapeutics for patients with cancer cachexia-induced heart disease. Current models of cancer cachexia-induced cardiac complications use rodents, or their cells, and although they offer valuable insights, they are limited owing to low throughput, short cell survival time ex-vivo and their restricted ability to recapitulate the chronic and complex nature of cardiac atrophy in humans (5, 14, 15). Thus, these approaches can fail to fully capture the progressive structural and functional changes in the heart caused by cachexia, affecting their capacity to test therapeutic interventions that promote cardiac recovery. In sum, to date, there is no cure for cancer cachexia-induced cardiomyopathy nor any effective treatments to prevent, halt or reverse disease (16–21).

It is clear that new models that employ human tissue for tracking cardiac impairment associated with cancer cachexia are needed. By harnessing advances in regenerative medicine, particularly through the use of human-induced pluripotent stem cell-derived cardiac muscle (hiPSC-CMs), new opportunities arise to overcome these limitations. HiPSC-CMs provide a unique system to model human-specific cardiac function in the context of cachexia. HiPSC-CMs have been highly useful in studies of human heart diseases (22–26). HiPSC-CMs can be used to create patient-specific cardiac models, enabling a more personalized understanding of chronic disease effects on cardiac health (25, 26). Furthermore, using hiPSC-CM models provides a platform to identify molecular mechanisms and to evaluate new therapeutic strategies aimed at mitigating cancer cachexia-induced cardiac dysfunction, making them ideal for testing the effects of novel treatments over extended periods.

In this study, we investigated the mechanism of cancer cachexia-induced cardiomyopathy using a new human iPSC-derived cardiac muscle experimental platform. We compared the use of transwell insert technology with cancer cell-conditioned medium to induce the cachectic phenotype. Data shows that cachectic hiPSC-CMs replicate the cardiac phenotype of cancer cachexia patients along multiple dimensions, including morphological, metabolic, and functional characteristics. Cachectic hiPSC-CMs enabled new mechanistic insights into the protein degradation pathways involved in the pathogenesis of cancer cachexia-induced cardiac dysfunction. Specifically, we report herein a central role of the Atrogin-1/calcineurin A/NFAT axis in mediating cancer-induced cardiac atrophy in this setting.

## Methods

### HiPSC Culture and Differentiation

For the human induced pluripotent stem cell line, we utilized the well characterized DF 19–9–11 line (Wicell) (27). HiPSC line was derived from newborn male foreskin dermal fibroblasts and is transgene and vector free, karyotype normal, and pluripotent, shown to have excellent capacity to differentiate into cardiac myocytes (28). hiPSCs were cultured as previously described (28); cells were grown in mTeSR media (Stemcell, Vancouver, CA) in plastic dishes coated with Matrigel (Corning, Corning, NY), passaged using gentle dissociation media when they reached 90% confluency, and replated with Rho kinase inhibitor 10 μM Y-27632 (Selleckchem, Houston, TX).

Human iPSCs were differentiated when they reached 70-95% confluency in 12 well plates according to the small molecule Matrigel sandwich method (29). Briefly, at day 0, cells were treated with Matrigel dissolved in RPMI + B27 supplement minus insulin (Thermo Fisher, Waltham, MA) with 8 μM CHIR99021 (Stemgent, Lexington, MA), which is a GSK3 inhibitor. On day 1, media was changed without the addition of small molecules. On day 2, cells were treated with 8 μM IWP-4 (Stemgent, Lexington, MA), an inhibitor of Wnt signaling, in RPMI + B27 supplement minus insulin. On day 4 and every 2 days after that, media was changed until cells started to beat vigorously, which is at approximately days 8–14, at which point media was switched to RPMI + B27 supplement with insulin. hiPSC-CMs underwent glucose starvation for cardiac myocyte purification as previously described (30), cardiac troponin T antibody (Thermofisher, MA5-12960-1:100) was used for flow cytometry studies. Media was replaced every 2 days. Media was supplemented with antibiotic-antimycotic (Thermo Fisher Scientific, Waltham, MA). For all the assays that involved dissociation and re-plating of hiPSC-CMs, cells were dissociated with accutase (Thermo Fisher, Waltham, MA) or trypsin at 37°C, followed by dissociation in 15% FBS + RPMI with a P1000 pipet and centrifugation for 3 minutes at 800 rpm.

### Rat ventricular myocyte isolation and primary culture

Animals used in these experiments were handled in accordance with guidelines set by institutional animal care and use committee (IACUC) and approved by University of Minnesota. Hearts were obtained from Sprague–Dawley rats. Adult rat ventricular myocyte isolation was performed as previously described (31, 32). Briefly, adult rats were anaesthetized by inhalation of isoflurane followed by i.p. injection of heparin (15,000 U/kg) and Fatal Plus (150 mg/kg). Following enzymatic digestion by retro-grade perfusion with collagenase and gentle trituration of the cardiac ventricles, cardiac myocytes were plated on laminin-coated glass coverslips (2 x 10^4^ myocytes/coverslip) and cultured in M199 media (Sigma, supplemented with 10 mmol/l glutathione, 26.2 mmol/l sodium bicarbonate, 0.02% bovine serum albumin, and 50 U/ml penicillin-streptomycin, with pH adjusted to 7.4, additionally insulin (5 μg/ml), transferrin (5 μg/ml) and selenite (5 ng/ml) (ITS) were added (Sigma I1884)). One hour after plating, nonadherent cells were removed and fresh M199 was applied.

### Tumor cell lines

Colon-26 (C26) mouse adenocarcinoma cells (Division of Cancer Treatment and Diagnosis Tumor Repository, National Cancer Institute), and HCT116 adult human male colorectal carcinoma cells (ATCC, CCL-247). Tumor cells were cultured in RPMI 1640+L-glutamine medium (Gibco, Thermo Fisher Scientific) supplemented with 5% (v/v) fetal bovine serum (FBS) and 1% (v/v) Penicillin–Streptomycin (10,000 U/ml) at 37°C with 5% CO2.

### Cachexia induction

Two protocols were used in inducing cachexia, either conditioned medium or co-culture of tumor cells with hiPSC-CMs using transwell inserts (CORNING, 353494), where insert membranes with 0.4 μM pores made of tissue culture treated-Polyethylene terephthalate (PET) are placed on the top of the iPSCs-CM culture plates (33).

Tumor cells were seeded in their growth media (5% FBS in RPMI) in the transwell inserts or plastic plates at 10-15% confluency and left for overnight attachment, the growth medium was removed the next day, and the cells were washed twice with sterile PBS and incubated in iPSCs – CM maintenance media (RPMI + B27+) to acclimatize the cells with the new media. For the supernatant (conditioned media) group: after 24 hours, the medium was collected and centrifuged at 5,000 rpm for 10 min, the collected media is used to condition media for cachexia induction (30% of the total iPSCs-CM maintenance media). For the transwell group, fresh media was changed the day of the experiment on the transwell and then the insert was co-cultured on the top of the iPSCs-CM to allow for gradual release of the tumor cells’ secretome in the media to mimic the tumor in-vivo kinetics. The control group was fed with fresh iPSCs maintenance media only. These steps were repeated daily for the duration of the cachexia induction. Three- and four-days protocols were followed for inducing cachexia with C26 and HCT116 cell lines respectively (33, 34).

Post-cachexia stage. Studying cachexia remission was done by changing media to fresh iPSCs-CM maintenance media for all groups, media changes are done every two days, and cells were tracked for one week in the remission stage.

A summary of the timeline and groups is shown in figures (1, 2 and 6).

#### Light microscopy imaging

Cells were imaged in 12 well plates using Keyence microscope for phase contrast imaging using 4X and 10X magnifications.

#### Metabolic flux analysis

Oxygen consumption rates (OCR) and extracellular acidification rates (ECAR) of hiPSC-CMs were measured using a Seahorse XFe96 Analyzer (Seahorse Bioscience, MA, USA). Cells were seeded onto matrigel-coated XFe96 96-well microplates at a denisty about 30,000 cells per well and left for one week until beating is observed again to ensure their functionality. Cachexia was induced in the Seahorse plate using the protocol described above. Prior to the assay, the media were removed and replaced by 180 μl of 2 mM glutamine, 10mM glucose and 1mM Sodium Pyruvate assay medium, without sodium bicarbonate, to allow for degassing the cells for OCR and ECAR measurements. The cells were preincubated for 1h at 37°C and 0% CO2 oven before loading into a Seahorse Bioscience XFe96 analyzer. During these 60 min, the ports of the cartridge containing the oxygen probes were loaded with the compounds to be injected during the assay and the cartridge was calibrated. ATP synthesis-linked O2 consumption and proton leak-driven respiration were determined by the addition of oligomycin (3.5 μM, Sigma, # O4876). After 3 measurement cycles, the uncoupler carbonilcyanide p – triflouromethoxyphenyl hydrazone (3 μM FCCP, Sigma, # C2920) was added to promote maximal respiratory capacity. After a further 3 measurement cycles, rotenone (3 μM, Sigma, R8875) was added to block complex I in addition to antimycin A (3 μM, Sigma, A8674) to inhibit complex III, thereby ablating mitochondrial oxygen consumption. Results were normalized to total protein concentrations.

#### Electrical stimulation and live-cell imaging

Electrical stimulation of monolayer cell cultures was performed using 12-well C-Dish (IonOptix, Westwood, MA) connected to a C-Pace EP Culture Pacer (IonOptix) applying a square, bimodal, biphasic voltage waveform with controllable amplitude, duration, and frequency.

Live-cell imaging was performed using a Nikon eclipse microscope (TE2000-U) in a stage-top physiological environmental chamber with 37°C temperature, 5% CO2, and humidity control (Ibidi, Germany), which was customized to fit the electrical pacing system. Real-time cardiac myocyte contraction images were recorded with an C13440 camera (ORCA-Flash4.0; Hamamatsu Photonics, Hamamatsu, Japan). Videos were analyzed using MUSCLEMOTION plugin [31] for FiJi Image J (35). Traces with the same pace only were included in the results.

### Western Blot and Protein Quantification

Cells were lysed with RIPA buffer (Sigma, # R0278) supplemented with cOmplete™ protease inhibitor (Roche, #11836153001) and PhosStop (Roche, #4906845001) on ice. Quantification was then done with a Pierce BCA kit (Thermo Fisher, Waltham, MA). 15-30 μg of non-boiled protein in Laemmli buffer with β-mercaptoethanol was loaded into each lane, and samples were run on 12 % precast polyacrylamide gels (BioRad) at 120 V until the dye front reached the end of the gel. Protein was transferred to PVDF membranes. Membranes were stained with primary antibodies (Atrogin-1 (ab168372, Abcam-1:1000), MuRF1 (55456-1-AP, Proteintech-1:1000), GAPDH (60004-1-Ig,

Proteintech-0.5:1000 or Ab37168, abcam-0.5:1000), Cardiac actin (AC1-20.4.2, Progen-1:1000), LC3B (#2775S, CellSignaling-1:300), Ubiquitin (80992-1-RR, Proteintech-1:1000 or Ab120 FK2, LifeSensors-1:1000), Calcineurin A (07-1492, Millipore sigma-1:1000 or 68163-1-IG, Thermofisher-1:1000) in TBS-T with 5% milk overnight at 4°C. Secondary antibodies were applied at a dilution of 1:10,000 for 1 hour in room temperature, and membranes were imaged on the LI-COR Odyssey (LI-COR, Lincoln, NE). Western blots were quantified using Image Studio Lite Version 5.2.5.

### Autophagy flux assessment

Bafilomycin A1 (CellSignaling, #54645) was used to treat control and cachectic hiPSC-CM (BAFA1, 200 nm) for 5 hours, followed by LC3B immunostaining using Western blot.

### Pro-Q diamond and Sypro ruby staining

Proteins separated by SDS-PAGE were visualized by in gel staining with Pro-Q Diamond (Invitrogen/Molecular Probes), a fluorescent dye specific for phospho-amino acids following the manufacturer’s protocol. Peppermint Stick Phosphoprotein molecular weight standards (Invitrogen/Molecular Probes), a molecular weight marker for detection of phosphorylated proteins was used as positive and negative control for phosphor-protein. Stained gels were visualized with ChemiDoc imager (Bio-rad). The same gel was then washed twice in distilled water and re-stained overnight with a solution of SYPRO Ruby dye for total proteins as per the manufacturer’s protocol. Then the gels were visualized with ChemiDoc imager (Bio-rad).

#### Immunofluorescence

hiPSCs were grown and differentiated on glass bottom plates (Cellvis, #P06-20-1.5-N and #P24-1.5H-N) into hiPSC-CMs for L/D and NFAT staining studies, except for cell size determination experiments where hiPSC-CMs were re-plated after differentiation to the glass bottom plates. Then cachexia was induced as mentioned earlier. hiPSC-CMs were fixed in a 4% paraformaldehyde solution for 10 minutes. Cells were permeabilized for 15 minutes with 0.1% triton. Prior to incubation with antibodies, hiPSC-CMs were blocked with 3% BSA in PBS. Conjugated Phalloidin antibody (Thermofisher, catalog # A12379) in blocking solution (diluted at 1:200) + DAPI (diluted at 1:500) was incubated 1 hour at RT and then washed off with PBS. This was followed by cell size determination in rat cardiac myocytes and hiPSC-CM. For Live/dead imaging: Calcein AM (Thermofisher, C3100MP), propidium iodide (Cayman chemical, 10008351) and Hoechst 33342 (CellSignaling, 4082) were used to stain live cells in culture. NFATC4 (Novus bio, NBP1-46210-1:50) was incubated for 1 hour at RT. All images were taken at 10X or 20X magnification using Nikon AX R confocal or Keyence fluorescence microscopy.

#### Cytokine Analysis

Cell lysates were tested by the Cytokine Reference Laboratory (CRL, University of Minnesota). This is a CLIA’88 licensed facility (license #24D0931212). Samples were analyzed on the Luminex platform using the mouse or human-specific multiplex cytokine/chemokine panel and the specific multiplex myokine panel on the Luminex platform. Samples were assayed for Luminex according to the manufacturer’s instructions. Fluorescent color-coded beads coated with a specific capture antibody were added to each sample. After incubation and washing, a biotinylated detection antibody was added followed by phycoerythrin-conjugated streptavidin. The beads were read on a Luminex instrument (Bioplex 200), a dual-laser fluidics-based instrument. One laser determines the analyte being detected via the color-coding; the other measures the magnitude of the PE signal from the detection antibody, which is proportional to the amount of analyte bound to the bead. Samples were run in duplicate, and values were interpolated from 5-parameter fitted standard curves.

### RNA isolation and gene expression analysis

Cells were solubilized in RLT buffer and RNA was extracted using the RNAeasy kit (Qiagen) and reverse transcribed using the iScript cDNA synthesis kit (Bio-rad) following the manufacturer’s recommendation. Gene expression analysis was carried out using 500 ng of cDNA per reaction in technical triplicates and qRT-PCR was performed using SYBR green master mix (PowerUp, Applied Biosystems) against: GAPDH (ENSMUSG00000057666): Forward CGT CCC GTA GAC AAA ATG GT, Reverse TTG ATG GCA ACA ATC TCC AC, IL6 (ENSMUSG00000025746): Forward CCA GAG TCC TTC AGA GAG ATA CA, Reverse AAT TGG ATG GTC TTG GTC CTT AG, TBP (ENSG00000112592): Forward GGT GCT AAA GTC AGA GCA GAA, Reverse CAA GGG TAC ATG AGA GCC ATT A, PUM1 (ENSG00000134644): Forward TGG ACC ATT TCG CCC TTT AG, Reverse CAG AGA GTT GTT GCC GTA GAA, IL6 (ENSG00000136244): Forward CCC TGA CCC AAC CAC AAA, Reverse GGA CTG CAG GAA CTC CTT AAA, GAPDH (ENSG00000111640): Forward GAC CAC TTT GTC AAG CTC ATTT C, Reverse CTC TCT TCC TCT TGT GCT CTT G. Real-time PCR was performed in accordance with Minimum Information for Publication of Quantitative Real-Time PCR Experiments guidelines (36). Product identity was confirmed by melting curve analysis.

### Image analysis

Image analysis was carried out using cell profiler for nuclei count and FIJI using custom macros for rat cardiac myocyte diameter and hiPSCs-CM cell size determination.

### Statistical analysis

Statistical tests were calculated in GraphPad Prism, version 10.2 (GraphPad Software). One-way ANOVA was used to compare more than two groups with one independent variable, followed by Tukey’s correction for multiple comparisons. Paired T-test was used for comparing the same well nuclei count before and after cachexia phase. P <0.05 was considered statistically significant. The GraphPad Prism 10 software was used for graphic representation.

## Results

### Animal-based cancer cachexia validates transwell insert co-culture model in cachexia induction

To advance establishing experimental parameters, as a prelude to enable hiPSC-CM studies, (14, 15), we used adult rat cardiac myocytes along with either the conventional protocol using cancer cell conditioned media or the transwell insert as a newly introduced approach in the context of cancer cachexia-induced cardiomyopathy. Two different colon cancer cell lines that are well-established for their potential to induce cachexia, namely C26 and HCT116 cancer cells, were employed for this approach (34, 37). Experimental groups include control group that receive regular cardiac maintenance media, supernatant group that receive cancer cell conditioned media and the transwell group that has the cancer cells seeded on the insert bottom with cardiac myocytes growing on the plate underneath (fig. 1b, S1a). In each condition, the medium was freshly changed every 24 h. Cachexia was confirmed by reduced cardiac myocyte diameter as a readout (fig. 1d). Both transwell and supernatant groups exhibited significant reduction in cardiac myocytes diameter compared to the control group. Cardiac myocyte diameter reduced by about 32.95%, 40.2%, 39.69% and 36.92% for C26 supernatant, C26 transwell, HCT116 supernatant and HCT116 transwell groups respectively (figure 1c and S1b). The duration of this experiment lasted two days and stopped owing to the observation of marked cell death predominantly in the cachectic groups that limited the study of extended cachexia duration or the study of cardiac myocytes after cachexia cessation (post-cachexia phase).

**Figure 1:**
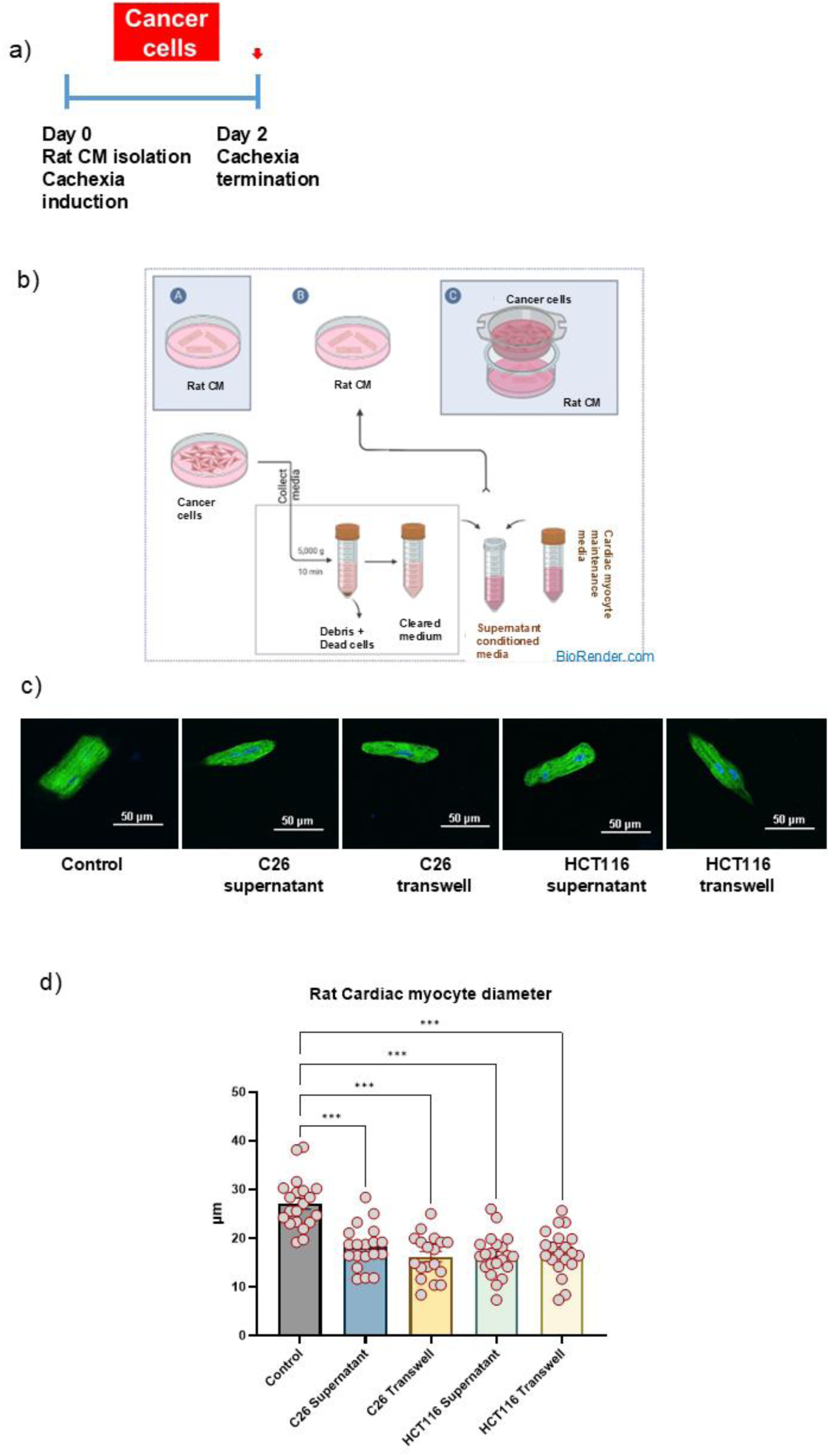
Cachexia induction in rat cardiac myocytes. a) Experimental timeline b) Experimental grouping for the model; [A) Control group receiving regular cardiac myocyte maintenance media, B) Supernatant group receiving cancer conditioned media & C) Transwell group, cells co-cultured with C26 or HCT116 cancer cell lines]. c) Representative image of the cachectic rat cardiac myocytes using C26 or HCT116 colon cancer cells, cells stained with phalloidin and DAPI and imaged using confocal microscopy. d) Reduction in diameter of cachectic rat cardiac myocytes (n=18-21 fiber/group). Data are presented as mean ± SEM, ANOVA test was performed to calculate P values; *P < 0.05, **P < 0.01, ***P < 0.001, and ****P < 0.0001.

The experimental timeline is shown in figure 1a. These findings help establish key methodological details but also highlight the limitation of the rodent cardiac myocyte system in studying cachexia over prolonged periods or in post-cachexia phase.

### Cachectic hiPSC-CM recapitulates the structural and morphological alterations of cancer cachexia-induced cardiac impairment

While the previously described cancer cachexia studies using rodents offer valuable insights into the pathology of cachexia-induced cardiac dysfunction (14, 15, 34, 37), this approach is limited by short survival time ex vivo. In addition, compared to in vivo studies, there is a need to understand the direct effect of cancer cachexia on the cardiac myocytes, which is difficult to achieve with bulk analysis of the heart due to the presence of non-cardiac myocytes. Therefore, we were motivated to establish a new experimental approach with increased throughput and longer survival to facilitate mechanistic study and form a platform for enabling future therapeutic testing. Thus, we generated an in vitro cancer cachexia induced cardiac impairment model using human cardiac myocytes generated from human induced pluripotent stem cells (hiPSCs).

Human cardiac muscle cells (hiPSC–CMs) were differentiated using human hiPSCs and purified using glucose free medium (Fig. S2, supplementary video S1). The cancer cachexia setup consists of either the pulse addition of conditioned media (supernatant group) or gradual release of cancer cell secretions (transwell group) as described above (Fig. 1a). The transwell mimics in vivo kinetics with the presence of the tumor cells in the system and the continuous release of their secretion in the media. The medium is refreshed every 24 hours for both treatments. The hiPSC–CMs cells were induced to develop cachexia using C26 cells (3 days protocol) or HCT116 cells (4 days protocol), as shown in the timeline in figure 2a. Length of induction period was supported by the amount of secreted IL6 as an established cachexokine from each cell line (Fig. 2f) (38–41), together with evidence from previous studies that used C26 and HCT116 for cachexia induction (33, 34).

**Figure 2:**
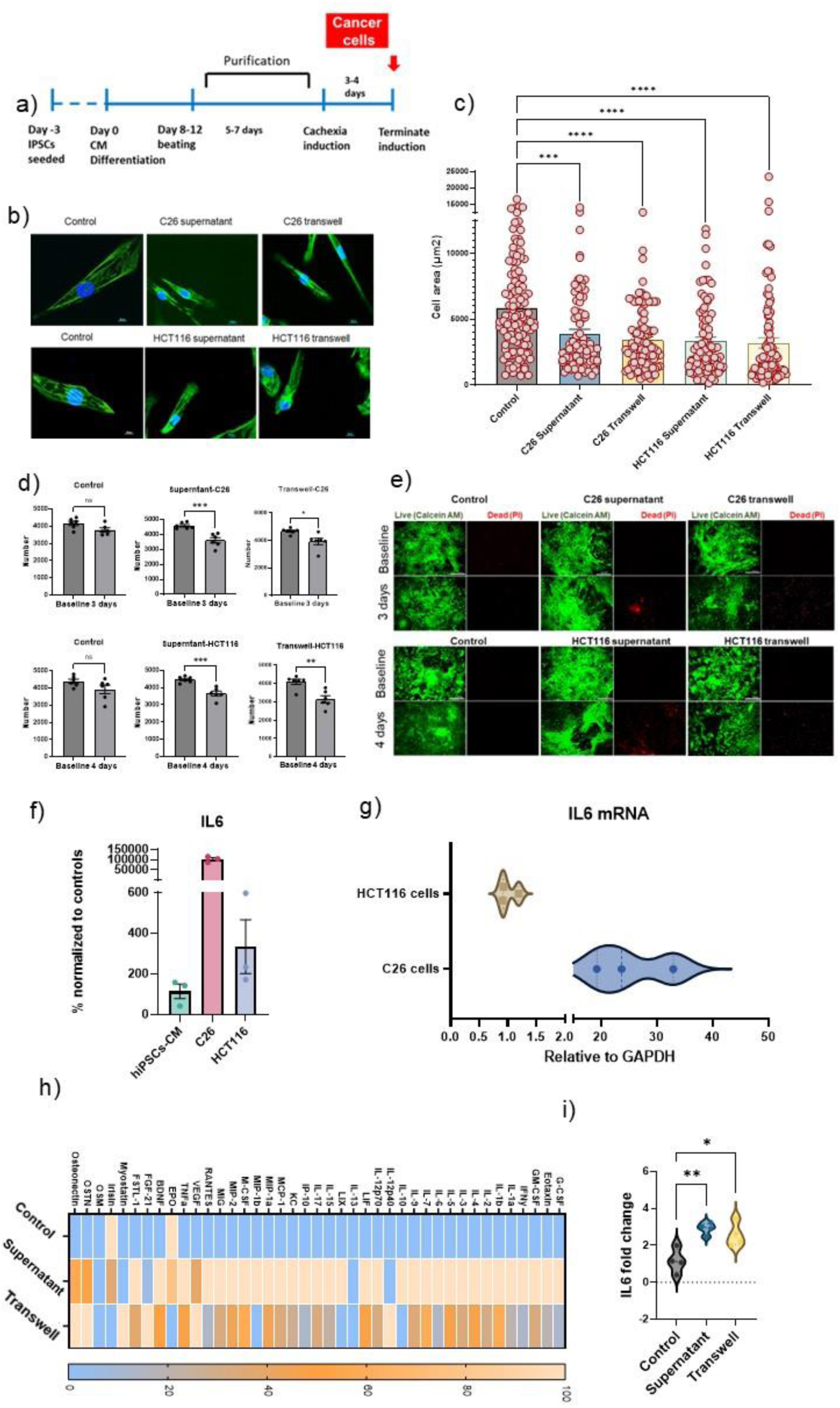
**Cachectic hiPSC-CMs, their morphological evaluation and inflammatory response.** a) Timeline of the experiment indicating cachexia phase. b) Cell size reduction in cachectic hiPSCs-CM as shown in the representative immunofluorescence images stained with Phalloidin and DAPI, Scale bar, 10 μm. c) Cell size reduction in cachectic hiPSC-CM (n=80-115/group). d) Nuclei number reduction in cachectic groups (supernatant and transwell) in the two tested models (C26 or HCT116), (n=6, paired T-test comparison). e) Representative images from control and cachectic hiPSCs-CMs showing cell death assessed using Live/dead cell viability assay. Live cells were detected with calcein AM (green) dye and dead cells were detected using propidium iodide (red). Scale bar, 300 μm. f) IL6 protein expression in the media of cultured cells; C26 or HCT116 cells compared to hiPSC-CM. g) qPCR analysis of IL6 mRNA expression in C26 and HCT116 cells. h) Cytokine array measured in the collected media at the end of cachexia phase from C26-CM cachectic model. Data normalized using Z-score analysis. i) qPCR analysis of IL6 mRNA expression in control and cachectic hiPSC-CM (C26-CM cachectic model). Data are presented as mean ± SEM, ANOVA test was performed to calculate P values; *P < 0.05, **P < 0.01, ***P < 0.001, and ****P < 0.0001.

The hiPSC-CMs developed cancer cachexia using supernatant treatment or transwell system, characterized by significant reduction in cardiac myocyte cell area, and decreased number of nuclei (Fig. 2b-d, S3a-f). Cell loss and dead cells as seen in live/dead imaging assay were observed in all the cachectic groups with prominent areas of cell death seen in the supernatant group that received conditioned media (Fig. 2e).

Both treatments with both the tested cell lines successfully induced cachexia in hiPSC-derived cardiac muscle, with cell area reduced by 33.45%, 41.46% for supernatant and transwell groups in the C26 treated group and by 42.82%, 45.51% for supernatant and transwell groups in the HCT116 treated group (Fig. 2c), indicating that both C26 and HCT116 colon cancer cell lines can induce cardiac wasting possibly through the release of pro-cachectic mediators that can cause cardiac atrophy.

Of note, with both cancer cell lines, there was a greater magnitude of atrophy when cardiac muscle was exposed to the transwell inserts containing cancer cells compared to supernatant group where cardiac muscle was exposed to conditioned media from cancer cells. On the other hand, decreased nuclei number and percentage of dead cells was more prominent in supernatant group compared to transwell group with both the tested cell lines.

### Pro-cachectic mediators driving the cardiac atrophy

As functions of interleukin-6 (IL6) contributing to cachexia pathogenesis have been well established (38–42), we characterized the expression profile of IL6 as a pro-cachectic mediator in the media collected from growing C26 or HCT116 tumor cell lines. IL6 protein levels showed significantly higher levels in both tumor cell lines compared to controls (hiPSC-CMs), with C26 cells showing higher expression levels compared to HCT116 cells (Fig. 2f). Similarly, IL6 mRNA levels were higher in C26 compared to HCT116 cells as measured by qPCR (Fig. 2g).

We next analyzed a cytokine/myokine array panel in the cultured media of cachectic groups, both supernatant and transwell groups, compared to control cardiac muscle group in the C26-CM model on day 3 (end of the cachexia phase). This profile reflects the crosstalk between cancer cell secretion and the pro-inflammatory response of cardiac myocytes during cancer cachexia phase. Cytokine profiling revealed a unique pattern in inflammatory response in the media collected from cachectic hiPSC-CMs (Fig. 2h). Several inflammatory cytokines, IL-1a, IL-1b, IFN-ɤ, IL-2, TNF-α, were upregulated in both cachectic groups compared to controls. Notably, IL-6 levels showed a 1000-fold higher in supernatant group (light orange) vs 150-fold higher in the transwell group (grey) compared to controls (blue) (Fig. 2h, S4a). Despite these quantitative differences in cytokine expression, the degree of cardiac atrophy observed was comparable in both conditions (Fig. 2b), suggesting that maximal atrophic signaling may occur even at lower cytokine thresholds. Additionally, IL6 mRNA levels were measured in the hiPSC-CMs at the end of the cachexia phase to assess the inflammatory response in the cachectic cardiac muscle compared to controls. Here, significant upregulation was detected in both cachectic groups with the supernatant group showing higher IL6 expression levels (Fig. 2i). This experiment thus identified a signature panel of cachexokines that are sufficient to drive cardiac myocyte atrophy in response to colon cancer growth independent from secondary host responses.

### Contractile and metabolic deficits in cachectic hiPSC-CMs

Cardiac dysfunction has been described as a severe complication in the cachectic state as induced by different cancer entities in both experimental animals and humans, including tumors of the colon, the lung, and the pancreas (8, 13, 14, 43–45). We thus paced cardiac cells to compare cells with the same rate, using C-pace IonOptix electrodes incubated in a gas, humidity and temperature-controlled chamber, and compared cardiac contractility between control and cachectic hiPSC-CMs using a video-based method as detailed above and illustrated in supplementary figure 5a-b. The contractility amplitude significantly decreased in the cachectic hiPSC-CMs (supernatant and transwell groups) compared to the control hiPSC-CMs. This trend was consistently observed among both tested models (C26-CM and HCT116-CM). In contrast, the relaxation time of the cachectic hiPSC-CM was significantly prolonged, with similar results in both models (Fig. 3b-g, supplementary videos S2-4). These results are evidence that in vitro cancer cachexia induction in hiPSC-CM can effectively replicate the contractile dysfunction observed clinically in cancer cachexia patients (44). It is worth mentioning that irregular beats along with resistance to follow the electrical pacing stimuli were noted in all cachectic groups, with a higher incidence in the transwell group of either the C26 or HCT-116 induced cachectic cardiac muscle (Fig. 3h, S5c).

**Figure 3:**
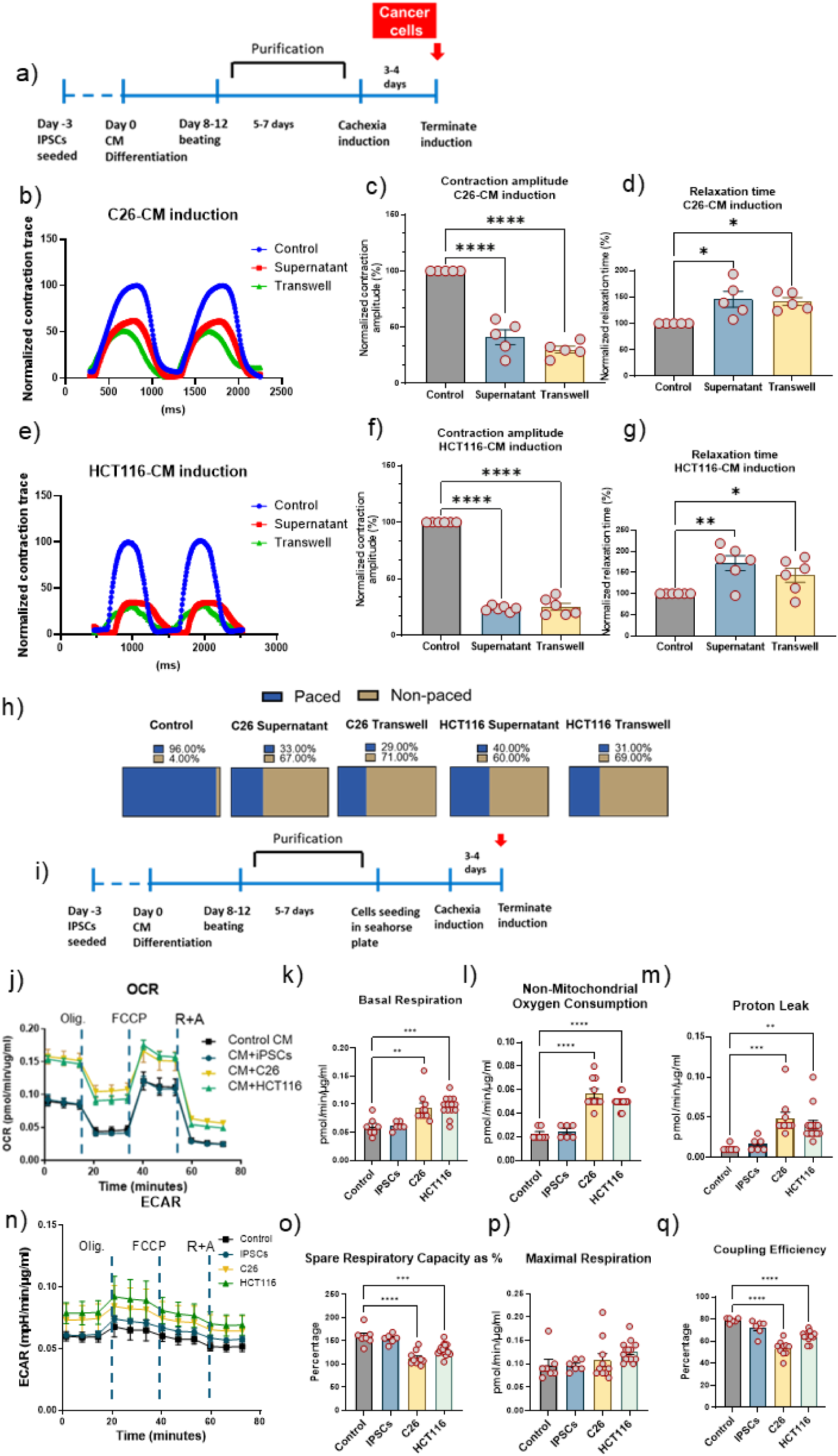
**Contractile function and metabolic profile of cachectic cardiac muscle.** a) Timeline of the experiment indicating cachexia phase. b) Representative trace visualizing paced contraction curves of C26 induced cachectic hiPSC-CM compared to controls. c-d) Quantitative analysis of hiPSC-CM contraction amplitude and relaxation time in C26 induced cachectic hiPSC-CM compared to controls. e) Representative trace visualizing paced contraction curves of HCT116 cachectic hiPSC-CM compared to controls. f-g) Quantitative analysis of hiPSC-CM contraction amplitude and relaxation time in HCT116 induced cachectic hiPSC-CM compared to controls. h) Proportion of arrhythmic hiPSC-CM in response to C-pace electrode stimulation across all groups. Recorded videos are analyzed using MUSCLEMOTION plugin and imageJ software. Traces not following pace or out of linearity were excluded from data analysis. i) Experimental timeline of the metabolic studies indicating cachexia phase. j-n) Representative mean kinetic profile of OCR and ECAR in cachectic hiPSC-CM (induced by conditioned media of C26 or HCT116 cells compared to control hiPSC-CM receiving regular cardiac muscle maintenance media only or conditioned media from growing hiPSCs. Quantification of (k) basal OCR, (l) non-mitochondrial oxygen consumption, (m) proton leak, (o) percentage spare respiratory capacity, (p) maximal respiration, and (q) coupling efficiency in cachectic and control hiPSC-CM. Data is presented as mean ± SEM; n=6-12 wells/group for each experiment. ANOVA test was used to calculate P values;; *P < 0.05, **P < 0.01, ***P < 0.001, and ****P < 0.001.

Given that metabolism plays a fundamental role in cancer cachexia and could potentially derive cardiac contractile impairment (41, 46–48), we next assessed the effect of cancer cachexia on the metabolic rates of the cachectic hiPSC-CMs. We used Seahorse technology to measure metabolic rates in the cachectic hiPSC-CMs and quantified various parameters. This experiment was done using conditioned media for cachexia induction, as the Seahorse plate is not compatible with the available transwell inserts in the market. We also used hiPSCs-conditioned media (collected from growing induced pluripotent stem cells) as an additional control group for this experiment to check if regular cell metabolites will change cardiac myocyte metabolic rates. We observed significantly higher basal respiration, non-mitochondrial oxygen consumption and proton leak levels in the cachectic hiPSC-CMs vs the control hiPSC-CMs. However, since maximal respiration was not significantly altered, spare respiratory capacity and coupling efficiency of the cachectic hiPSC-CMs were significantly reduced compared to controls (Fig. 3j-q). Data from two cachectic models showed a consistent trend and altogether suggested an energy wasting condition associated with higher metabolic rates with inefficient energy utilization. To evaluate whether these increased basal respiration levels in the cachectic hiPSC-CMs can be attributed to cachectic proteins released from the cancer cells or due to altered metabolites, we repeated the Seahorse experiment after boiling supernatants prior to media conditioning to denature proteins released from the cancer cells. Boiling the media abrogated the observed increase in basal respiration levels, resulting in comparable oxygen consumption rates between the cachectic and control hiPSC-CMs (Fig. S6a-b).

### LC3B and FBXO32/Atrogin-1, but not MuRF1, underlies atrophic remodeling in cachectic hiPSCs-derived cardiac muscle

To gain molecular insight into the mechanisms of cachexia development and cardiac myocyte wasting in hiPSC-CMs, we investigated the two main protein degradative pathways involved in cardiac proteostasis, autophagy and the ubiquitin-proteasome system (UPS) (8, 41, 49). LC3BII/I levels are used as a major marker of autophagic activity (44, 50, 51), and an indicator of autophagosome abundance (51). We measured the LC3BII/I ratio to determine whether autophagy-mediated protein degradation contributes to cardiac wasting in cancer cachexia. Our results showed a marked reduction of LC3B-I levels in cachectic hiPSC-CMs compared to control hiPSC-CMs across the two tested models (Fig. 4b-c), suggesting LC3B-I consumption to form LC3B-II. However, LC3B-II levels remained comparable among all the investigated groups. Calculating the LC3B-II/I ratio showed a significant increase in the cachectic hiPSC-CMs (Fig. 4b-c). However, since the LC3B-II protein levels did not rise among the tested groups, we sought to investigate autophagy flux to determine whether the lack of elevation in LC3B-II levels was due to increased autophagy flux or other upstream defects in this signaling pathway. Autophagic flux represents the dynamic process including autophagosome generation, fusion with lysosomes, and the rapid degradation of autophagic substrates within autolysosomes along with LC3B-II. Autophagic inhibitors such as Bafilomycin A1 impede the degradation of autolysosome contents by inhibiting the Na+/H+ pump in the lysosome, leading to increased lysosomal pH and inhibition of acidic lysosomal proteases. This leads to an accumulation of autophagosome structures (51). Upon using such inhibitors, accumulation of LC3B-II demonstrates efficient autophagic flux, while in case of a failure of LC3-II protein to increase under the inhibitor effect, it indicates a defect or delay earlier in the autophagic process, prior to degradation at the autolysosome. To investigate this, we treated control and cachectic hiPSC-CMs with Bafilomycin (BAFA1, 200 nm) for 5 hours. This leads to accumulation of LC3B-II confirming that cachectic cells exhibit increased autophagy flux (Fig. 4d). Collectively, our analysis of the LC3B immunoblots demonstrated higher LC3B-I levels in control hiPSC-CMs suggesting slower lipidation, a process required for active autophagy execution. In contrast, cachectic hiPSC-CMs showed decreased LC3B-I levels indicating that LC3B-I is being lipidated faster, with an active autophagy flux. Together, these findings indicate elevated autophagy execution in the cachectic hiPSC-CMs, suggesting that autophagy is a major contributor to human cardiac muscle atrophy in cancer cachexia.

**Figure 4:**
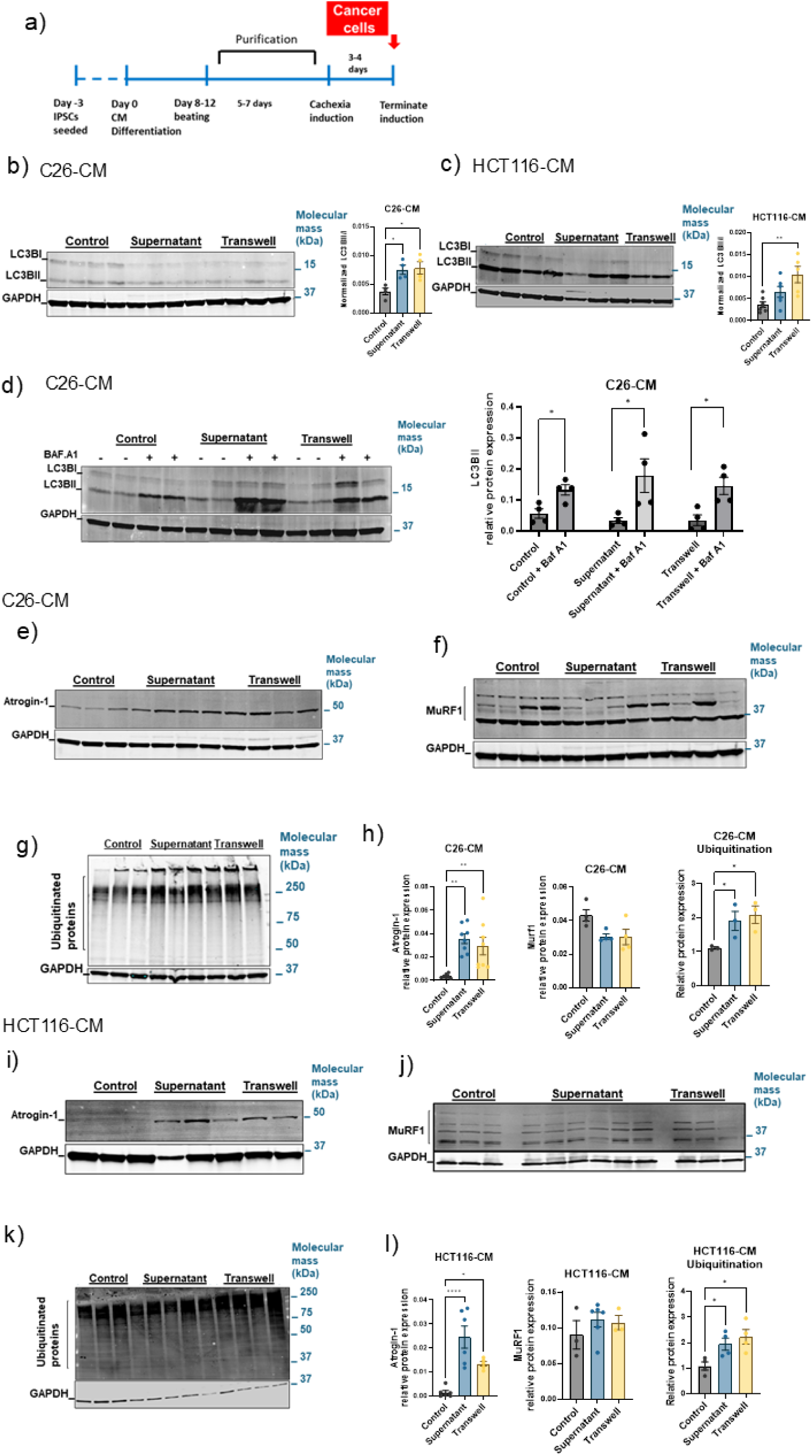
Protein degradation markers in control and cachectic hiPSC-CM. a) Timeline of the experiment indicating cachexia phase. LC3BI/II levels in C26 cachectic (b) and HCT116 cachectic (c) models. LC3BI/II levels after BafA1 treatment in C26 cachectic model (d). Immunoblot analysis for Atrogin-1 in C26 cachectic (e) and HCT116 cachectic (i) models. Murf-1 in C26 cachectic (f) and HCT116 cachectic (j) models. Global ubiquitinated proteins in C26 cachectic (g) and HCT116 cachectic (k) models. h,l) Quantified Atrogin-1, MuRF1 protein levels and ubiquitinated proteins in C26 or HCT116 cachectic models, normalized to GAPDH. Data is presented as mean ± SEM, experiments done in triplicates. ANOVA test was performed to calculate P values; *P < 0.05, **P < 0.01, ***P < 0.001, and ****P < 0.0001.

We next examined the ubiquitin E3 ligases FBXO32/Atrogin-1 and TRIM63/MuRF1 as a striated muscle-specific UPS markers and critical regulators of skeletal muscle size and mass. Atrogin-1 and MuRF1 are well documented to be upregulated in cancer cachexia induced skeletal muscle atrophy (52–54). Nonetheless, while protein degradation pathways in skeletal muscle and in the heart are thought to be the same, it remains unclear whether they are regulated in the same manner, particularly in human tissue (55). Therefore, we examined whether cancer cachexia induction in hiPSC-CMs would lead to an increase in Atrogin-1 and MuRF1 expression. Following cachexia induction, the expression of Atrogin-1 was elevated in the hiPSC-CMs in both the supernatant and transwell groups (Fig. 4e,i). On the contrary, MuRF1 proteins levels remain unchanged between groups (Fig. 4f,j). Furthermore, the levels of ubiquitinated proteins demonstrated modest but statistically significant increases in cachectic hiPSC-CMs compared to control hiPSC-CMs (Fig. 4g,k). Data from the two models (HCT116 and C26) showed a consistent result. These data suggest that enhanced UPS-mediated protein degradation is contributing to the observed cardiac muscle atrophy in the cachectic hiPSC-CMs besides increased levels of Atrogin-1 does not markedly affect the global UPS function, rather we speculate that it specifically affects the turnover of specific substrates.

Overall, these results demonstrate that the autophagy effector LC3B and the ubiquitin ligase FBXO32/Atrogin-1 are important regulators of protein degradation during cachectic atrophic remodeling of the heart.

### Calcineurin A (CnA) – NFAT Pathway in the atrophic cardiac muscle

We next tested whether Atrogin-1 upregulation induces cardiac atrophy in response to cancer cachexia and plays a role in the cardiac cell size reduction and contractile dysfunction. Among targets of atrogin-1 in striated muscle, calcineurin A (CnA), a calcium/calmodulin dependent serine/threonine protein phosphatase is prominent because of its role in coordinating cardiac myocyte gene expression programs that determine cell size in the context of cardiac hypertrophy (55–57). To investigate this axis in cardiac atrophy, we measured α-calcineurin (CnAα) levels among the different experimental groups. Strikingly, α-calcineurin levels show a distinct pattern in the cachectic groups of the tested models. We observed a significant reduction of calcineurin at 32 kDa MW band which is indicative of the active constitutive form of α-calcineurin (Fig. 5b-d). This form has been previously reported in various cell types and represents the

**Figure 5:**
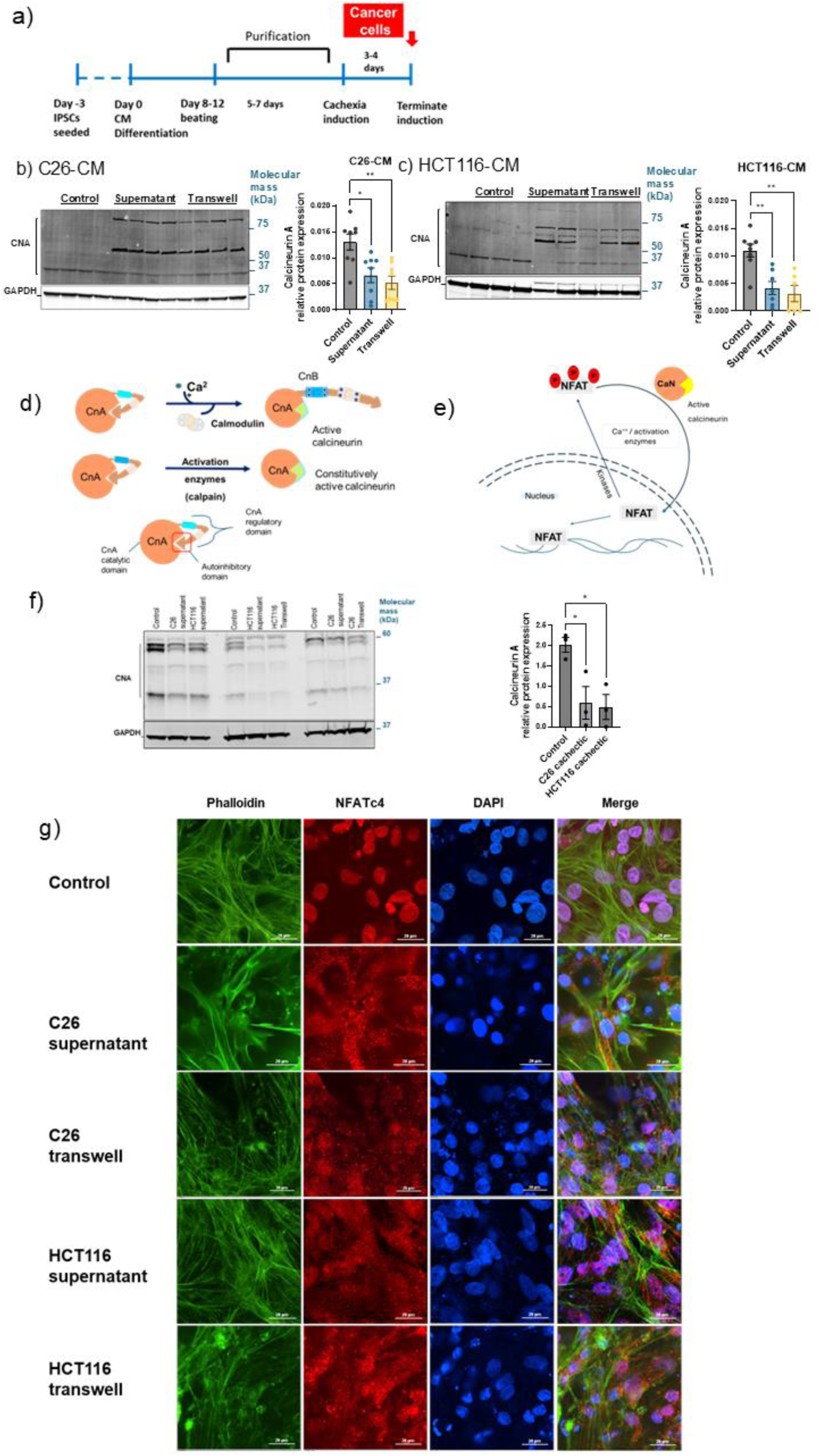
**Calcineurin axis studies**. a) Timeline of the experiment indicating cachexia phase. Immunoblot analysis of α-Calcineurin (32 kDa) levels in C26-CM (b) and HCT116-CM (c) models. d) Schematic illustration of calcineurin structure, activation and the constitutive active form of calcineurin. e) Schematic illustration for calcineurin phosphatase activity and NFAT downstream effect. f) Immunoblot analysis of α-Calcineurin (60 kDa) levels using antibody specific to N-terminal of α-Calcineurin. g) Immunofluorescence confocal imaging for control and cachectic hiPSC-CMs. Cells stained with DAPI, phalloidin and NFATc4. Scale is 20 μm. Data is presented as mean ± SEM, experiments done in triplicates. ANOVA test was performed to calculate P values; *P < 0.05, **P < 0.01, ***P < 0.001, and ****P < 0.0001.

catalytic functional domain of the catalytic subunit (58–61). In addition, we observed high MW bands exclusively in cachectic groups, suggesting a modified form of the protein that resulted from cachexia induction (Fig. 5b,c). Together with the upregulated Atrogin-1 (E3 ligase), this may indicate ongoing ubiquitination of α-calcineurin (62). Further we utilized an additional CnAα antibody that targets the N-terminal region of CnAα, detecting the full length CnAα (60 kDa). Notably, the full length CnAα exhibited a significant reduction in protein levels compared to control hiPSC-CMs (Fig. 5f). Based on these results, we proposed that Atrogin-1 induces cardiac atrophy in response to cancer cachexia via degradation of CnA. Additionally, reduced calcineurin Aα levels could underlie the observed decreased cardiac myocyte size and the contractile dysfunction possibly through the downstream NFAT signaling.

### NFAT signaling in cachectic hiPSC-CMs

Given that cancer cachexia disrupted the Atrogin-1/CnA axis in the cardiac muscle, we speculated that this might affect the CnA–NFAT pathway in the cachectic cardiac muscle. Calcineurin A directly activates nuclear factor of activated T-cells (NFAT) nuclear translocation and transcriptional activity (63, 64). Nuclear translocation of NFAT occurs rapidly after dephosphorylation by calcineurin A and is required for its transcriptional activity (Fig. 5e). To this end, we evaluated NFAT protein localization and expression by immunostaining at the end of the cachexia phase with an antibody against NFATc4 (NFAT3). Nuclei were counterstained with DAPI (Fig. 5g). Cachexia induction leads to increased cytoplasmic localization of NFATc4 in the cachectic hiPSC-CMs across both models. In addition, the nuclear staining of NFAT increased in control hiPSC-CMs (Fig. 5g), suggesting the effect of Atrogin-1 on CnA activity and cardiac atrophy is cell-autonomous. Taken together, these results suggest that the upregulated Atrogin-1 can affect NFAT signaling via calcineurin regulation in the context of cardiac atrophy.

Given that calcineurin is a calcium/calmodulin dependent serine/threonine protein phosphatase, we tested the effect of dysregulated Atrogin-1/CnA axis on cardiac muscle protein phosphorylation. We used the pro-Q diamond global phosphorylated protein probe with SYPRO ruby total protein stain to evaluate differential phosphorylation in cachectic vs control hiPSC-CMs. Our results indicated that the decreased calcineurin levels contributed to a decrease in phosphatase activity in the cachectic cells as evidenced by the increased phosphorylation of protein band within 110-170 kDa range (Fig. S7a-d). This band corresponds to the molecular weight of NFAT proteins (65), suggesting that the dysregulated CnA protein level impacts its activity on specific protein targets rather than exerting a global effect. The results provide additional evidence that calcineurin A plays a significant role in the pathogenesis of the cachectic cardiac muscle, which ultimately leads to cardiac atrophy in response to cancer cachexia.

We have investigated another potential Atrogin-1 target, alpha actin (66), but in contrast to CnA, the protein levels of alpha actin were not affected in cachectic hiPSC-CMs in both C26 and HCT116 models (Fig. S8a-b).

### Post-cachexia studies

As an extension of this work, we evaluated the hiPSC-CMs ability to recover from muscle wasting and cachexia in the post-cachexia phase. We removed the conditioned media and the insert with the tumor cells growing and changed media freshly to regular cardiac maintenance media for one week. Morphological measurements done on single cell seeded hiPSC-CMs at the end of the one-week period revealed that cells recovered from significant size reduction, and cell sizes were not significantly smaller relative to controls (Fig. 6b-c).

**Figure 6:**
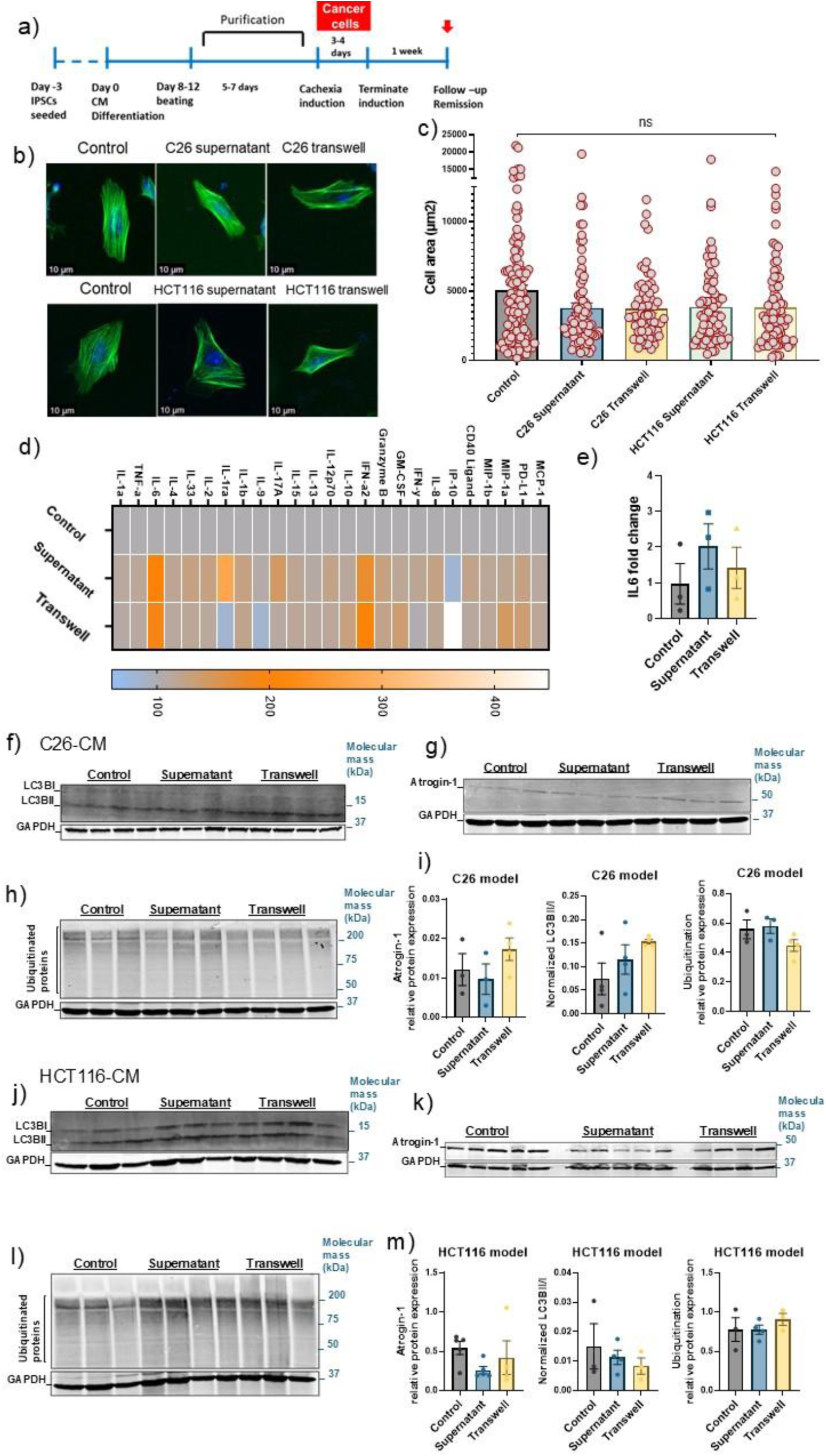
**Post-cachexia studies:Morphological, inflammatory and protein degradation.** a) Timeline of the experiment indicating post-cachexia phase. b) Cell size comparison in post-cachectic hiPSC-CM as shown in the representative immunofluorescence images stained with Phalloidin and DAPI, Scale bar, 10 μm. c) Cell size of post-cachectic hiPSC-CM (n=70-140/group), red line indicating the trend of the cell size change. d) Cytokine profile measured in the collected media at the post-cachexia phase (C26-CM model). e) qPCR analysis of IL6 mRNA expression post-cachectic hiPSC-CMs (C26 and HCT116 models). Data normalized as percentage comparison to controls. Immunoblot analysis for the LC3BI/II levels (f), Atrogin-1 (g), and global ubiquitinated proteins (h) in C26 post-cachectic model, values are quantified in (i), normalized to GAPDH. Immunoblot analysis for the LC3BI/II levels (j), Atrogin-1 (k), and global ubiquitinated proteins (l) in HCT116 post-cachectic model, values are quantified in (m), normalized to GAPDH. Data is presented as mean ± SEM, n=3-4 samples/group. ANOVA test was performed to calculate P values; *P < 0.05, **P < 0.01, ***P < 0.001, and ****P < 0.0001.

Similar to the cachexia phase, media collected from cultured cardiac muscle in the post-cachexia phase at one week time point after cachexia cessation in the C26-CM model were used to assess levels of a cytokine/myokine array. Cytokines levels mostly returned to the baseline after the one-week time point, nevertheless IL6 levels retained modestly but non-significantly elevated levels compared to the controls at the one-week remission time point (Fig. 6d-e). However, these IL6 levels were markedly decreased compared to IL6 levels during the cachexia phase (Fig. S4b).

Importantly, to assess if any sequalae of disrupted protein degradation remains at the molecular level, we evaluated the hiPSC-CMs in the post-cachexia phase. LC3B-II/LC3B-I ratio among the cachectic groups was comparable to the control levels (Fig. 6f,j). Atrogin-1 protein levels were normalized as compared to controls (Fig. 6g,k). Similar to the results observed during the cachexia stage, MuRF1 levels showed no significant differences (Fig. S9). Moreover, global ubiquitinated protein levels exhibited no significant differences across the tested groups in both models (Fig. 6h,l). Altogether, these results indicate cachexia remission of the hiPSC-CMs at the protein degradation molecular level. Tracking the post-cachectic hiPSC-CMs after Atrogin-1 levels have returned to normal levels revealed no significant difference between CnA levels among control and post-cachectic cardiac muscle (Fig. 7b,c).

**Figure 7:**
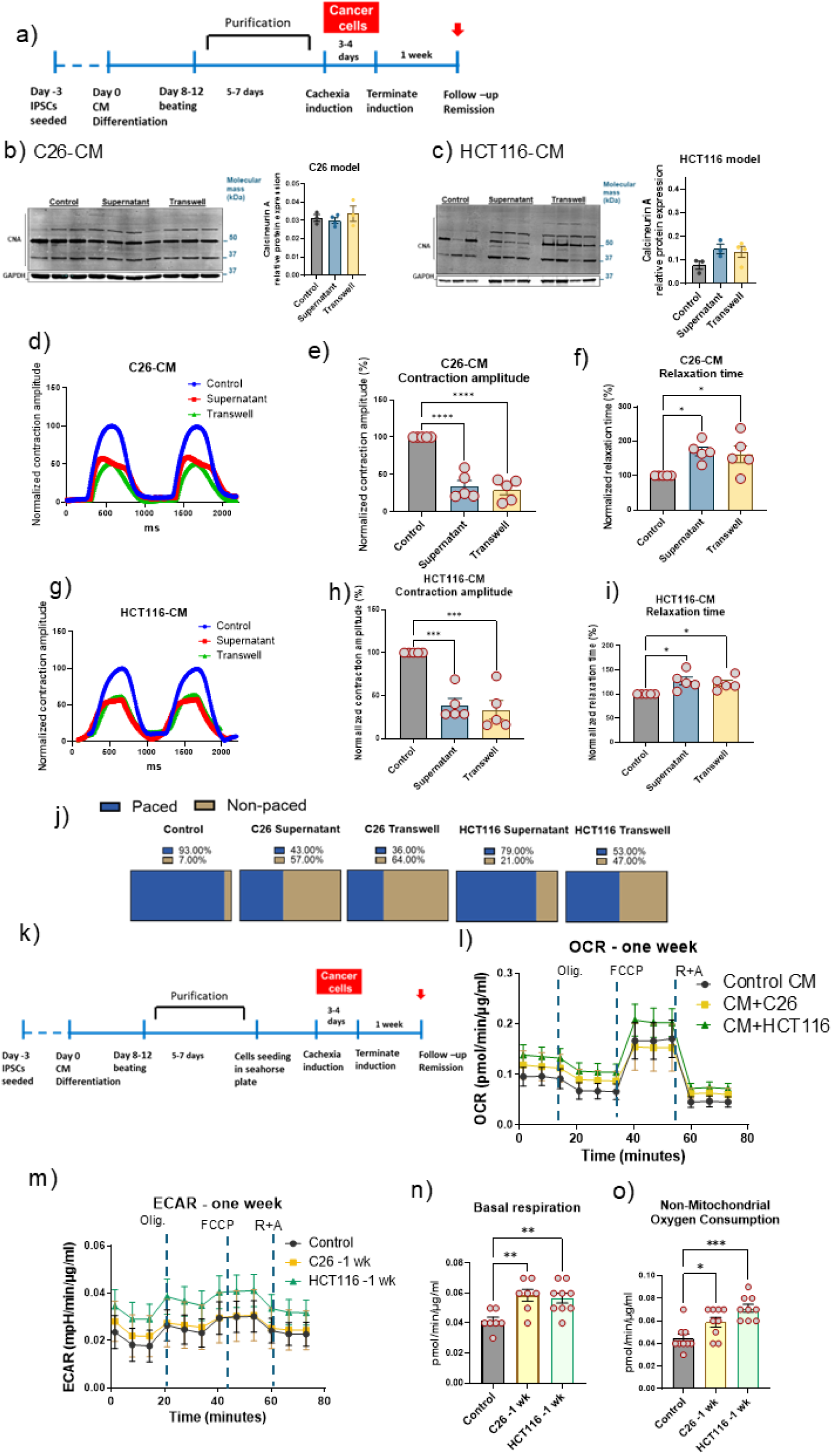
**Post-cachexia studies: Molecular, functional and metabolic.** a) Timeline of the experiment indicating post-cachexia phase. Immunoblot analysis of α-calcineurin protein levels for C26-CM (b) and HCT116-CM (c) post-cachectic models, with their quantified values normalized to GAPDH. d) Representative trace visualizing paced contraction curves of C26-CM model in post-cachexia stage compared to controls. e-f) Quantitative analysis of hiPSC-CM contraction amplitude and relaxation time in C26-CM model in post-cachexia stage. g) Representative trace visualizing paced contraction curves of HCT116-CM model in post-cachexia stage compared to controls. h-i) Quantitative analysis of hiPSC-CM contraction amplitude and relaxation time in HCT116-CM model in post-cachexia stage compared to controls. j) Proportion of arrhythmic hiPSC-CM in response to C-pace electrode stimulation across all groups. Recorded videos are analyzed using MUSCLEMOTION plugin and imageJ software. Traces not following pace or out of linearity were excluded from data analysis. k) Experimental timeline of the metabolic studies indicating post-cachexia phase. l-m) Mean kinetic profile of OCR and ECAR in C26 or HCT116 –CMs models at the one-week time point. Quantification of (n) basal OCR and (o) non-mitochondrial oxygen consumption for the hiPSC-CM in C26 or HCT116 –CMs models at the one-week time point. Data is presented as mean ± SEM; experiments done in triplicates. ANOVA test was used to calculate P values; *P < 0.05, **P < 0.01, ***P < 0.001, and ****P < 0.001.

The metabolic and functional insult observed in the cachectic cardiac muscle further prompted us to evaluate the cells in the post-cachexia phase at the one-week time point, considering the clinical characteristics of cancer survivors to bench mark our model (44, 67). The functional deficit persisted (Fig. 7d-j), reflecting the residual impaired cardiac function observed in cachectic patients even after cancer removal, which contributes to decreased survival in this population (44, 67). Regarding the metabolic phenotype, the upregulated basal respiration persisted even after the removal of the conditioned media with its secreted proteins and cachectic factors and the regular media change using the cardiac maintenance media for one week (Fig. 7l-o). This suggests that cachexia induction induced a sustained metabolic dysregulation that might be contributing to the long-term cardiac dysfunction.

## Discussion

We investigated the mechanism of tumor-initiated cardiac wasting-based cardiomyopathy using a novel hiPSC-CM cancer cachexia induction/remission system. Here, hiPSC-CMs effectively recapitulated the cancer cachexia-induced cardiac wasting cardiomyopathy observed in humans (44). The main findings of this study include elucidating a direct effect of tumor cells to induce cardiac wasting, all in the absence of cardiotoxic treatments. This distinction is important as anti-tumor treatments are themselves well known to cause cardiac disease leaving the tumor-dependent component unclear (68, 69). Specifically, our data show cardiac wasting involves increases in autophagy flux, via increases in LC3BII/LC3bI together with increases in E3 ligase Atrogin-1-mediated ubiquitination. Mechanistically, a new cardiac wasting signaling pathway emerges wherein we propose that the increased Atrogin-1 leads to degradation in Calcineurin A that, in turn, prevents the nuclear localization of the cardiac growth transcriptional regulator NFAT. We reasoned that the cytoplasmic retention of NFAT, caused by the cancer cachexia milieu, and in the absence of cardiotoxic interventions, can account for cardiac muscle wasting, as well as contributing to the consequent metabolic and contractile dysfunction, which are the hallmarks of cancer-mediated cardiomyopathy.

Our cardiac wasting findings reveal a complex interplay wherein both the autophagy and UPS pathways are implicated, with Atrogin-1, but not MuRF1, emerging as a critical player in the UPS pathway within the cancer cachexia cardiac wasting context. Atrogin-1 has been widely studied in skeletal muscle, where it regulates muscle atrophy by targeting Myo-D, a key transcription factor that is absent in cardiac tissue (66). However, in the heart, and only under the conditions of cardiac hypertrophy, has Atrogin-1 been previously shown to target calcineurin A (56). Our new findings provide new insights into Atrogin-1/calcineurin A axis and its role in modulating molecular mechanisms of cardiac atrophy in cachectic human iPSC-CMs.

Notably, this study extends previous research on calcineurin A that primarily focuses on cardiac hypertrophy, whereas here we present data supporting calcineurin A’s involvement in the pathological progression of cardiac atrophy. Calcineurin A is a calcium-dependent phosphatase and a known regulator of NFAT activity. Calcineurin A action entails the dephosphorylation of NFAT, promoting its nuclear translocation to induce gene expression leading to cardiac muscle growth (56, 70, 71). In the cachectic cardiac muscle, we observed that the altered calcineurin A levels markedly impaired nuclear translocation of NFAT, suggesting disrupted signaling limiting NFAT transcriptional activity. These data suggest that cardiac atrophy and altered gene expression observed in the cancer-induced cardiomyopathy conditions are caused at least in part by this NFAT altered localization and hence transcriptional dysregulation. It is noteworthy that the NFAT transcription factor acts synergistically with other transcription factors as NFkB, AP-1, GATA-4 and MEF-2 to assemble higher order transcriptional complexes affecting cardiac remodeling (72). Given the established role of NFkB in cancer cachexia induced skeletal muscle wasting (73–75), these interconnected regulatory nodes present new avenues for studying transcriptional factors contributing to skeletal and/or cardiac muscle wasting. This highlights the broader contribution of Atrogin-1, calcineurin A, and NFAT in cardiac remodeling. Another finding that fits well with hypertrophy regression, the recently introduced concept that involves the reversal of cardiac hypertrophy (76). Hypertrophy regression shares several common signaling pathways with cardiac atrophy but lacks evidence on Atrogin-1/Calcineurin A/NFAT axis involvement, a gap we address here.

Our data indicate that autophagic flux is elevated in the setting of cancer cachexia-induced cardiomyopathy. We observed a decreased level of LC3BI, however LC3BII exhibited comparable levels to controls, resulting in an elevated LC3BII/I ratio. This suggests either a block in autophagic degradation or increased autophagy flux. To test this, we treated cardiac myocytes with Bafilomycin A1, a lysosomal inhibitor. The resulting build-up of LC3BII upon Bafilomycin A1 treatment confirms that autophagosomes are actively formed under cachectic conditions, indicating an increase in autophagic flux, consistent with previous cancer cachexia-induced cardiomyopathy studies (7, 77, 78). Although autophagy is critical for maintaining cellular homeostasis, persistently elevated autophagy can culminate in cardiac cell loss (79). Thus, these findings support that excessive or dysregulated autophagy contributes to cardiac muscle wasting in cancer cachexia.

Importantly, the concurrent activation of both the autophagy-lysosome and Atrogin-1 ubiquitin E3 ligase indicates a coordinated action of proteolytic/degradative systems in cachectic hiPSC-CMs. While autophagy and UPS are essential for cellular quality control, their persistent or dysregulated activation in cancer cachexia likely contributes to pathological cardiac atrophy and functional decline. These findings suggest that targeting these proteolytic pathways may offer therapeutic potential in mitigating cardiac muscle loss in cancer cachexia.

This study provides other important contributions to the field. For example, a notable advance is that we employed the transwell system in the context of cancer cachexia-induced cardiac impairment, enabling paracrine interaction with cancer cells, further enhances the fidelity of the cachexia simulation. Our study confirmed the effectiveness of both approaches (conditioned media and transwell system) in modelling cachexia. Importantly, the transwell system, where cancer cells are co-cultured in proximity without direct contact, appeared to induce slightly more pronounced cell size reduction and greater degree of non-responsive hiPSC-CMs to pacing stimuli and irregular beats, suggesting that the presence of cancer cells in close proximity may exacerbate cachexia and functional disruptions. These findings suggest that cell-cell communication within the tumor microenvironment may involve complex signaling mechanisms beyond soluble factors, potentially including exosome-mediated transfer or direct membrane interactions.

Additionally, we observed an increase in cell loss within the supernatant groups where we used conditioned media to induce cachexia. This higher degree of cell death in the conditioned media-treated cardiac muscle may have impacted the experimental outcomes by potentially skewing functional assessments, particularly with respect to the remaining viable cell population. This observation is particularly important as it suggests that the conditioned media system, while inducing cachexia, may inadvertently introduce confounding variables due to the elevated cachexokine levels and elevated cell death, which could mask subtle phenotypic changes. On the other side, although the transwell system showed slightly more pronounced cell size reduction, it appeared to preserve a larger proportion of cells, offering a more stable platform. Thus, while both systems successfully induce cachexia, the transwell setup offers distinct advantages in terms of reducing excessive cell loss and providing a clearer picture of atrophic remodeling.

Additional contributions here reside in the fact that most cancer cachexia rodent studies show a decrease in metabolic rates in both skeletal and cardiac muscle (80, 81), which stands in contrast to the clinical presentation of cachexia patients (82–84), who generally have higher metabolic rates. Cachectic hiPSC-CMs closely replicates the human clinical phenotype, wherein we observed significantly higher oxygen consumption rates (OCR) in the cachectic cardiac muscle. This finding is consistent with a study on cachectic rat cardiac myocytes (15), suggesting that elevated metabolic activity may be a characteristic feature of cachectic cardiac myocytes. This increased OCR points towards the possibility of a “futile metabolic cycle,” wherein energy is consumed without being effectively utilized, potentially contributing to the rapid muscle wasting seen in cachexia. This metabolic dysregulation may help explain the heightened energy expenditure in cachectic patients, offering insights into the pathophysiological mechanisms underlying cardiac muscle metabolism during cachexia which eventually call for additional investigations. Cachectic hiPSC-CMs, therefore, offer a valuable platform for studying cachexia metabolic consequences, and aligns more closely with the clinical presentation of cachexia patients.

Another key advancement of using hiPSC-CMs is that it not only captured the morphological, functional, and metabolic phenotypes of human cancer cachexia-induced cardiomyopathy, but also extended to the post-cachexia phase. This provides the opportunity to investigate mechanisms of remission, recovery or any residual sequalae or deficit following cachectic insult. This is important as this area is largely underexplored in the context of cardiac muscle biology.

In summary, our new findings point to tumor-initiated alterations in the Atrogin-1/calcineurin A/NFAT signaling axis as contributor for cancer cachexia-induced cardiac wasting-based cardiomyopathy. This new finding is established in the absence of effects due to cardiotoxic treatments. Accordingly, these results provide new avenues for future therapeutic interventions to significantly improve patient outcomes and reduce the cardiovascular burden on cancer patients and survivors.

## Acknowledgment

We thank members of the Metzger lab for support and assistance. We acknowledge using resources of the University of Minnesota University Imaging Center (UIC). We gratefully acknowledge funding from NIH (HL132874, AR079477), AHA, Regenerative Medicine Minnesota (1184847) and UMN in support of this study.

