## Supplemental figures for "Elucidating cancer cachexia-mediated aberrant cardiac wasting signaling in human iPSC-derived cardiac muscle"

#### Supplementary figures legend

Fig.1: a) Experimental grouping for the model; [(A) Control group receiving regular cardiac myocyte maintenance media, B) Supernatant group receiving cancer conditioned media & C) Transwell group, cells co-cultured with C26 or HCT116 cancer cell lines]. b) Representative image of the cachectic rat cardiomyocytes using C26 or HCT116 colon cancer cells, cells imaged using Phase contrast - Keyence microscopy.

Fig.2: Purity of cardiac myocytes generated from hiPSCs. Percentage of cTNT+ cardiac myocytes using flow cytometry compared to unstained cells, before (a) and after b) purification.

Fig.3: a) Cell size comparison in cachectic hiPSCs-CM as shown in the representative immunofluorescence images stained with Phalloidin and DAPI, Scale bar, 20 μm. b) Images for cell size determination in cachectic hiPSCs-CM stained with Phalloidin and DAPI, Scale bar, 200 μm. c) Phase contrast representative image showing cachectic hiPSC-CM compared to control hiPSC-CM in C26-CM or HCT116-CM model, Scale bar, 100 μm. d) Images for cell size determination in post-cachectic hiPSCs-CM stained with Phalloidin and DAPI, Scale bar, 200 μm. e) Phase contrast representative image showing post-cachectic hiPSC-CM compared to control hiPSC-CM in C26-CM or HCT116-CM model, Scale bar, 100 μm. f) Example of morphometric measurement for cell size determination using Fiji software, Scale bar, 500 μm.

Fig.4: Raw values for IL6 protein levels measured in the collected media form cachectic (a) and post-cachectic (b) hiPSC-CM compared to controls in C26-CM model. Data is presented as mean ± SEM, n=4-9 samples/group. ANOVA test was performed to calculate P values; *P < 0.05, **P < 0.01, ***P < 0.001, and ****P < 0.0001.

Fig.5: a) Schematic figure for cardiac muscle contraction assessment. b) Photos of the functional studies set-up with its components: i-Closed temperature, humidity and gas-controlled chamber enclosing the plate of hiPSC-CM with the C-pace electrode inserted in the plate. ii-Closed ibidi chamber. iii-Top and bottom of the C-pace electrode, 12 well format with two electrodes for each well. iv-Ibidi temperature (top) or gas mixer (bottom) controllers. v-Ionoptix C-pace cell culture stimulator. c) The recorded video is subjected to analysis using musclemotion plugin and Fiji, example output traces are shown demonstrating traces responding to electric stimulus (paced) and non-respondents traces (non-paced).

Fig.6: a) Mean kinetic profile of OCR in hiPSC-CM treated by boiled conditioned media of C26 or HCT116 cells. Quantification of (b) basal OCR and (c) non-mitochondrial oxygen consumption in treated hiPSC-CM.

Fig.7: Protein visualization in HCT116-CM cachexia model. Proteins separated by SDS-PAGE were visualized by staining with a) Pro-Q Diamond (Invitrogen/Molecular Probes), a fluorescent dye specific for phospho-amino acids. As a molecular weight marker and as positive and negative control for detection of phosphorylated proteins we used Peppermint Stick Phosphoprotein molecular weight standards (Invitrogen/Molecular Probes) that contain a mixture of phosphorylated and non-phosphorylated proteins. The same gel was re-stained overnight with a solution of b) SYPRO Ruby dye for total proteins with their quantification.

c) Protein visualization in C26-CM cachexia model. Proteins separated by SDS-PAGE were visualized by staining with a) Pro-Q Diamond (Invitrogen/Molecular Probes), a fluorescent dye specific for phospho-amino acids. As a molecular weight marker and as positive and negative control for detection of phosphorylated proteins we used Peppermint Stick Phosphoprotein molecular weight standards (Invitrogen/Molecular Probes) that contain a mixture of phosphorylated and non-phosphorylated proteins. The same gel was re-stained overnight with a solution of d) SYPRO Ruby dye for total proteins with their quantification.

Fig. 8: Immunoblot analysis for α-actin in C26 cachectic (a) and HCT116 cachectic (b) models.

Fig. 9: Immunoblot analysis for MuRF1 in HCT116 post-cachectic model.

Fig. 10. Unedited whole blots:

- Immunoblot analysis for Atrogin-1 in C26 cachectic model.
- Immunoblot analysis for Atrogin-1 in C26 cachectic model-blot2.
- Immunoblot analysis for MuRF1 in C26 cachectic model.
- Immunoblot analysis for MuRF1 in C26 cachectic model followed by LC3B stain.
- Immunoblot analysis for global ubiquitination in C26 cachectic model.
- Immunoblot analysis for LC3B after BAFA1 treatment in C26 cachectic model.
- Immunoblot analysis for LC3B after BAFA1 treatment in C26 cachectic model-blot 2.
- Immunoblot analysis for calcineurin in C26 cachectic model.
- Immunoblot analysis for calcineurin in C26 cachectic model-blot2.
- Immunoblot analysis for calcineurin in C26 cachectic model-blot3.
- Immunoblot analysis for Atrogin-1 in HCT116 cachectic model.
- Immunoblot analysis for Atrogin-1, LC3B in HCT116 cachectic model-blot2.
- Immunoblot analysis for global ubiquitination in HCT116 cachectic model.
- Immunoblot analysis for calcineurin and LC3B in HCT116 cachectic model.
- Immunoblot analysis for calcineurin in HCT116 cachectic model.
- Immunoblot analysis for MuRF1 in HCT116 cachectic model.
- Immunoblot analysis for CNAα in C26 and HCT116 cachectic models.

- Immunoblot analysis for Atrogin-1 followed by global ubiquitination staining in C26 post-cachectic model.
- Immunoblot analysis for LC3B and calcineurin in C26 post-cachectic model.
- Immunoblot analysis for calcineurin in C26 post-cachectic model.
- Immunoblot analysis for Atrogin-1 in HCT116 post-cachectic model.
- Immunoblot analysis for global ubiquitination followed by LC3B stain in HCT116 post-cachectic model.
- Immunoblot analysis for calcineurin in HCT116 post-cachectic model.

a)

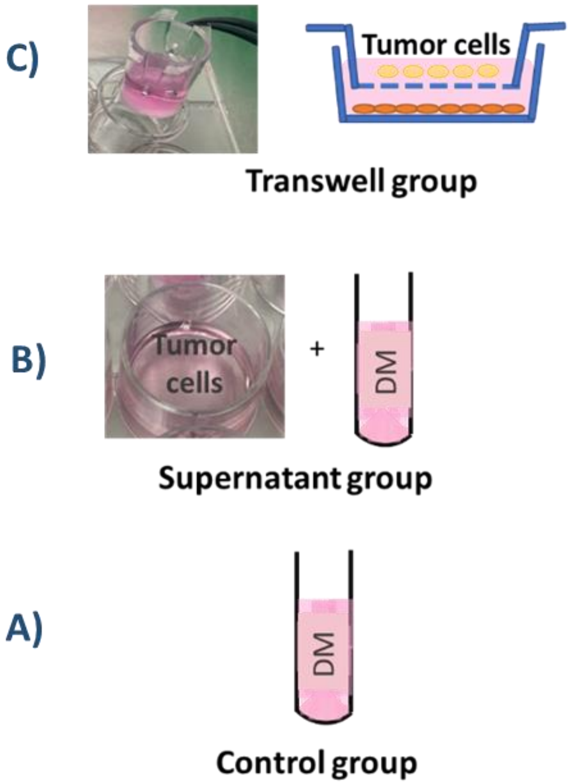

C)
Tumor cells
Transwell group
B)
Tumor cells
+
DM
Supernatant group
A)
DM
Control group


b)

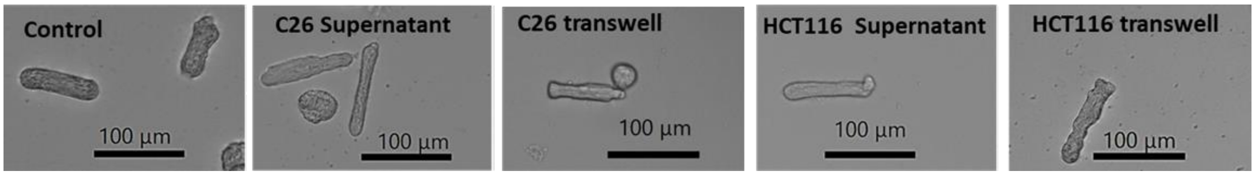

Control
100 µm
C26 Supernatant
100 µm
C26 transwell
100 µm
HCT116 Supernatant
100 µm
HCT116 transwell
100 µm


Figure 1.

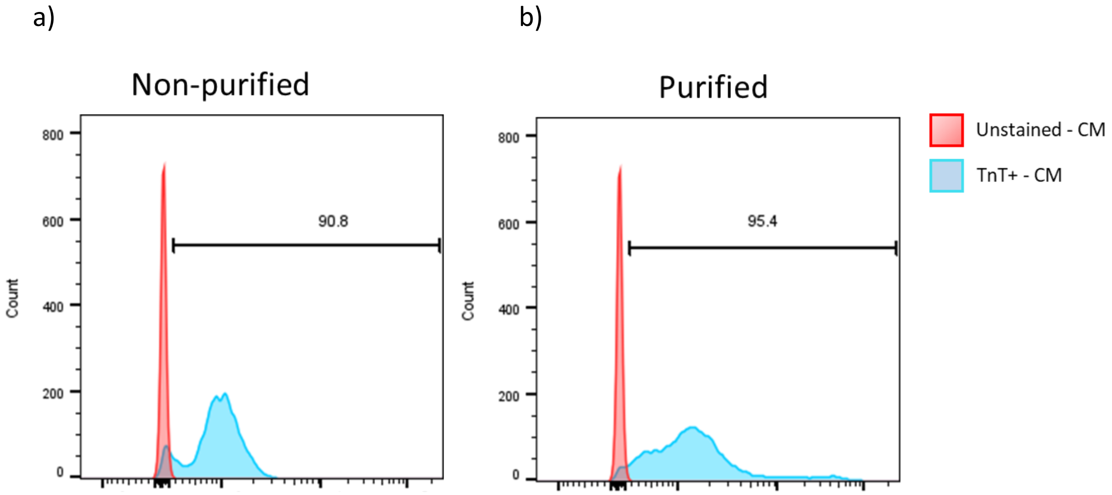
a)
b)
Non-purified
Purified
Count
800
600
400
200
0
90.8
95.4
Unstained - CM
TnT+ - CM

Figure 2.

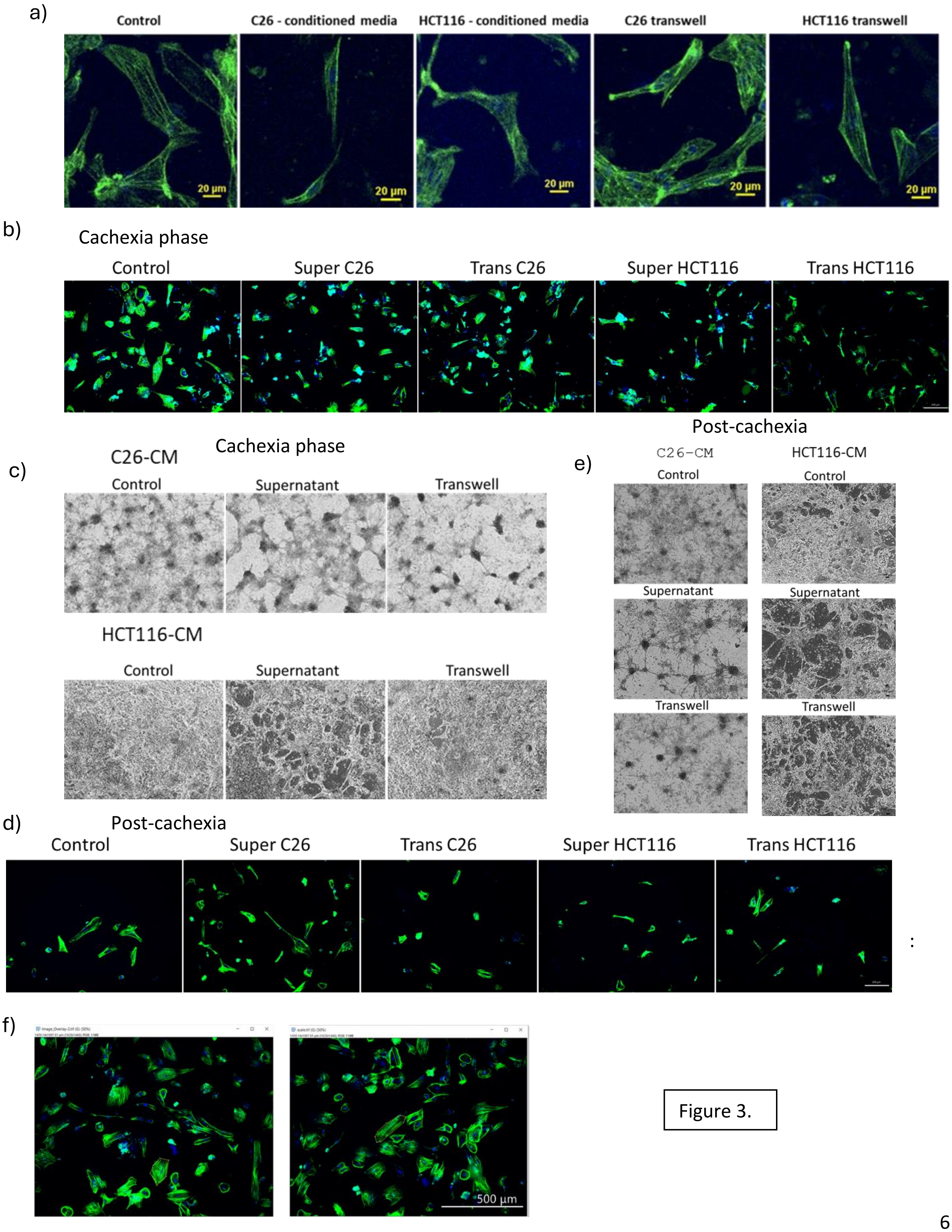
a)
Control
C26 - conditioned media
HCT116 - conditioned media
C26 transwell
HCT116 transwell
20 µm
b)
Cachexia phase
Control
Super C26
Trans C26
Super HCT116
Trans HCT116
c)
C26-CM
Cachexia phase
Control
Supernatant
Transwell
HCT116-CM
Control
Supernatant
Transwell
Post-cachexia
e)
C26-CM
HCT116-CM
Control
Control
Supernatant
Supernatant
Transwell
Transwell
d)
Post-cachexia
Control
Super C26
Trans C26
Super HCT116
Trans HCT116
f)
500 µm

Figure 3.

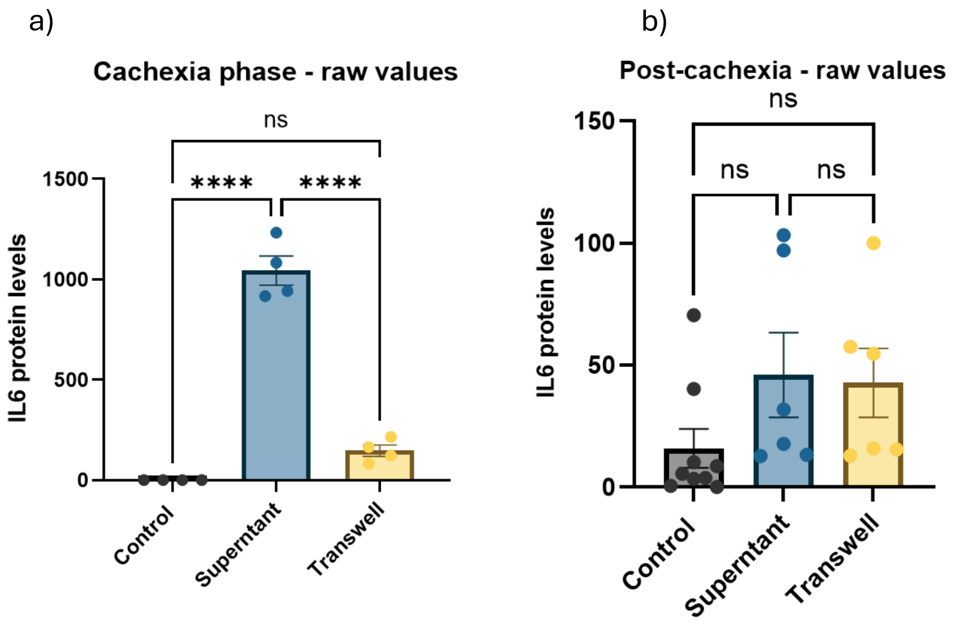
a)
Cachexia phase - raw values
ns
****
****
1500
1000
500
0
IL6 protein levels
Control
Supertant
Transwell
b)
Post-cachexia - raw values
ns
ns
ns
150
100
50
0
IL6 protein levels
Control
Superntant
Transwell

Figure 4.

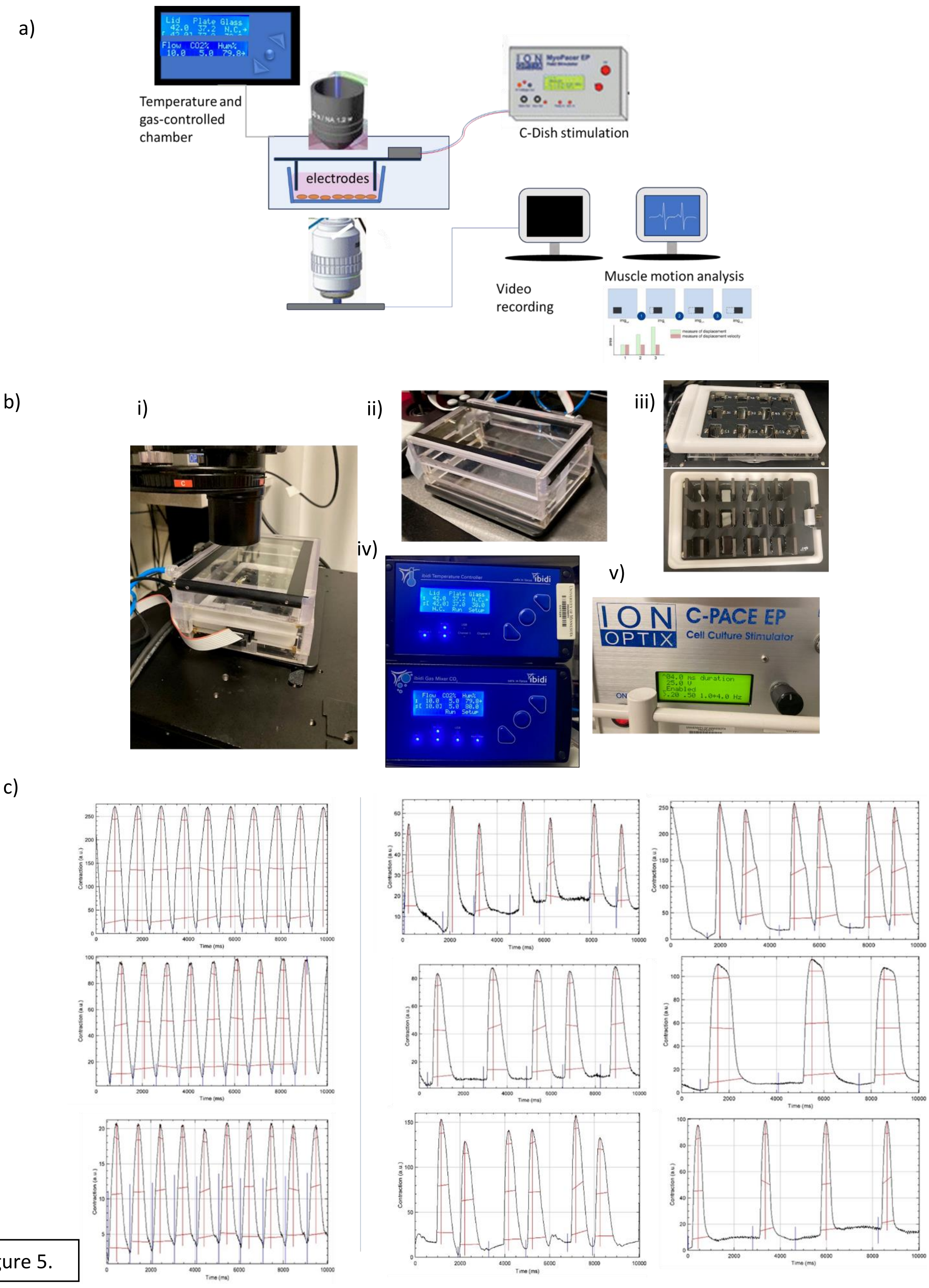
a)
Temperature and gas-controlled chamber
electrodes
C-Dish stimulation
Video recording
Muscle motion analysis
b)
i)
ii)
iii)
iv)
v)
ION OPTIX
C-PACE EP
Cell Culture Stimulator
c)
Contraction (a.u.)
Time (ms)

Figure 5.

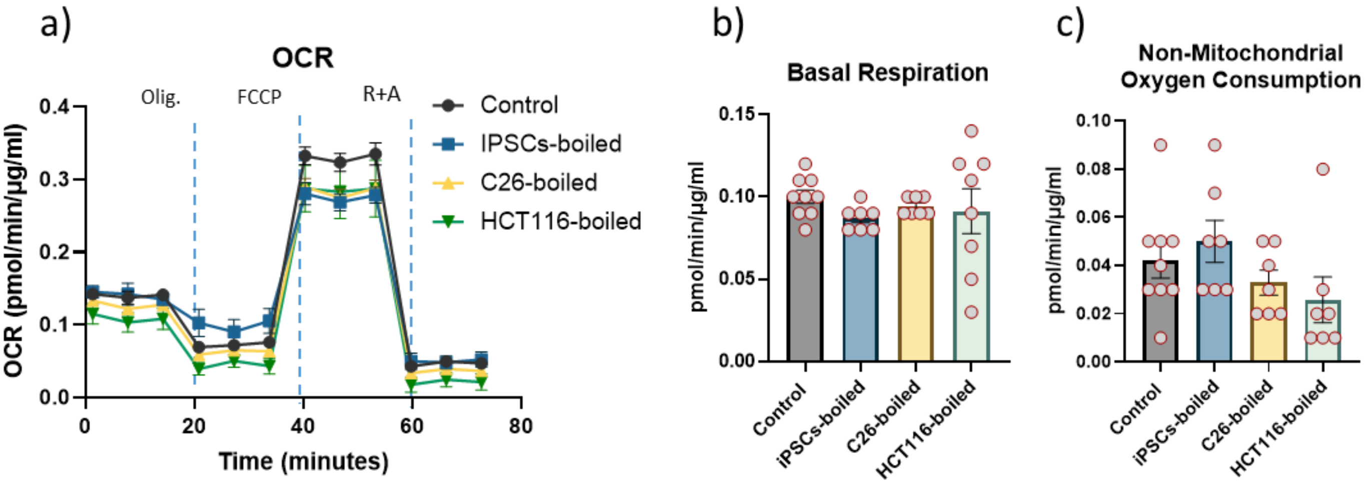

a)
OCR
Olig.
FCCP
R+A
Control
IPSCs-boiled
C26-boiled
HCT116-boiled
OCR (pmol/min/µg/ml)
Time (minutes)
b)
Basal Respiration
pmol/min/µg/ml
Control
iPSCs-boiled
C26-boiled
HCT116-boiled
c)
Non-Mitochondrial Oxygen Consumption
pmol/min/µg/ml
Control
iPSCs-boiled
C26-boiled
HCT116-boiled


Figure 6.

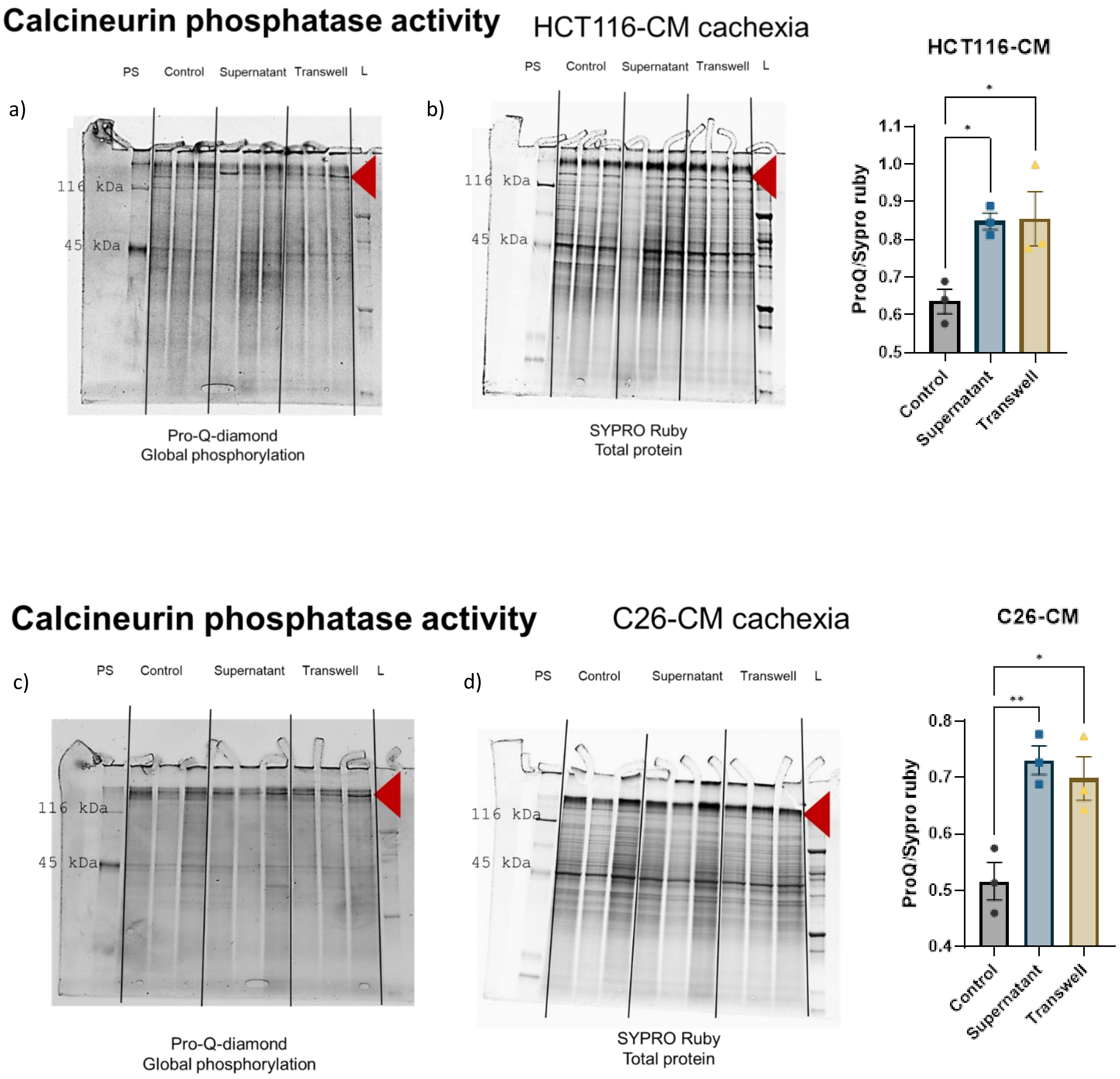
Calcineurin phosphatase activity
HCT116-CM cachexia
a)
PS Control Supernatant Transwell L
116 kDa
45 kDa
Pro-Q-diamond
Global phosphorylation
b)
PS Control Supernatant Transwell L
116 kDa
45 kDa
SYPRO Ruby
Total protein
HCT116-CM
ProQ/Sypro ruby
1.1
1.0
0.9
0.8
0.7
0.6
0.5
*
*
Control
Supernatant
Transwell
Calcineurin phosphatase activity
C26-CM cachexia
c)
PS Control Supernatant Transwell L
116 kDa
45 kDa
Pro-Q-diamond
Global phosphorylation
d)
PS Control Supernatant Transwell L
116 kDa
45 kDa
SYPRO Ruby
Total protein
C26-CM
ProQ/Sypro ruby
0.8
0.7
0.6
0.5
0.4
*
**
Control
Supernatant
Transwell

Figure 7.

a) C26-CM cachexia-alpha actin

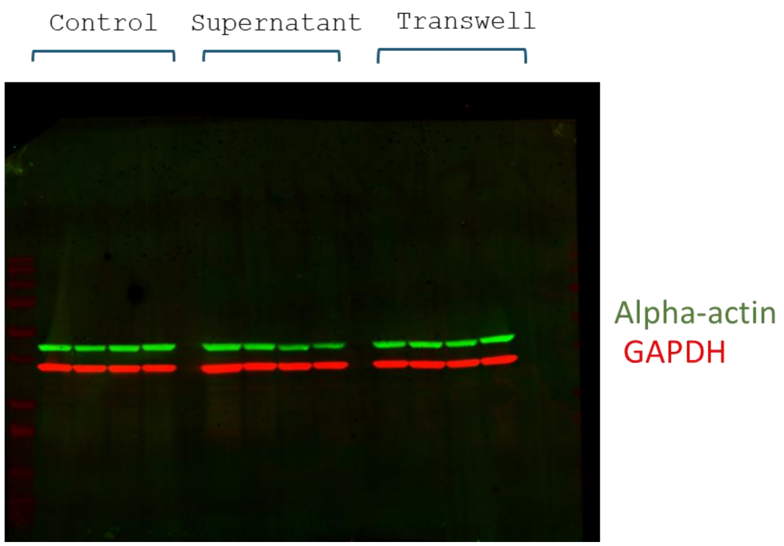


b) HCT116-CM cachexia – alpha actin

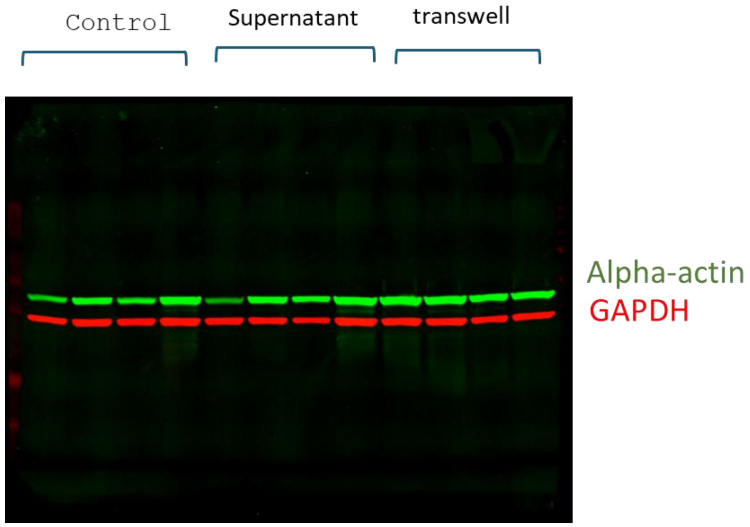


Figure 8.

**HC116-CM 1week**

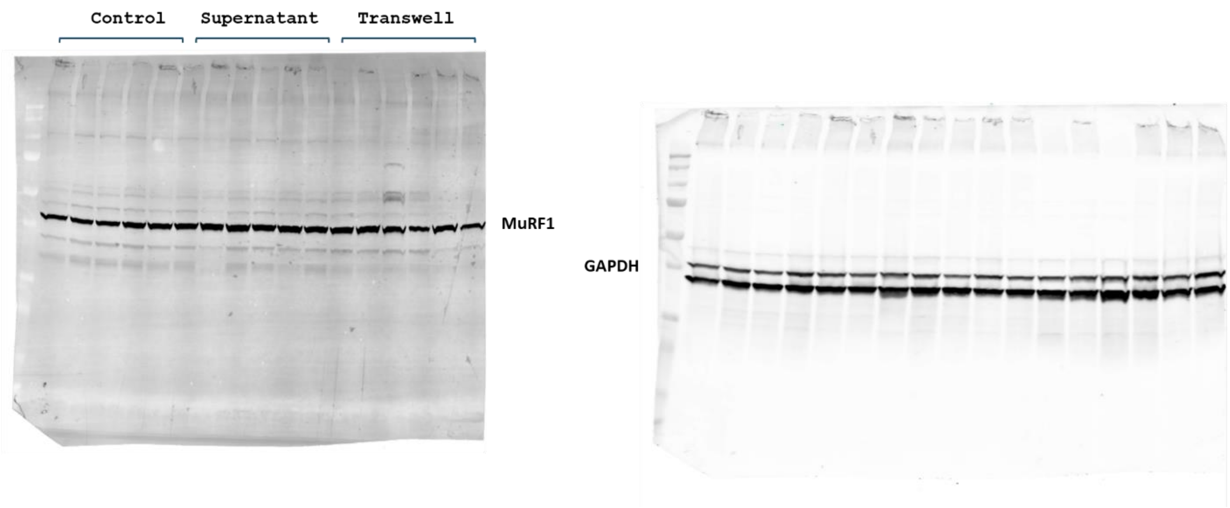


Figure 9.

Whole blots C26-CM

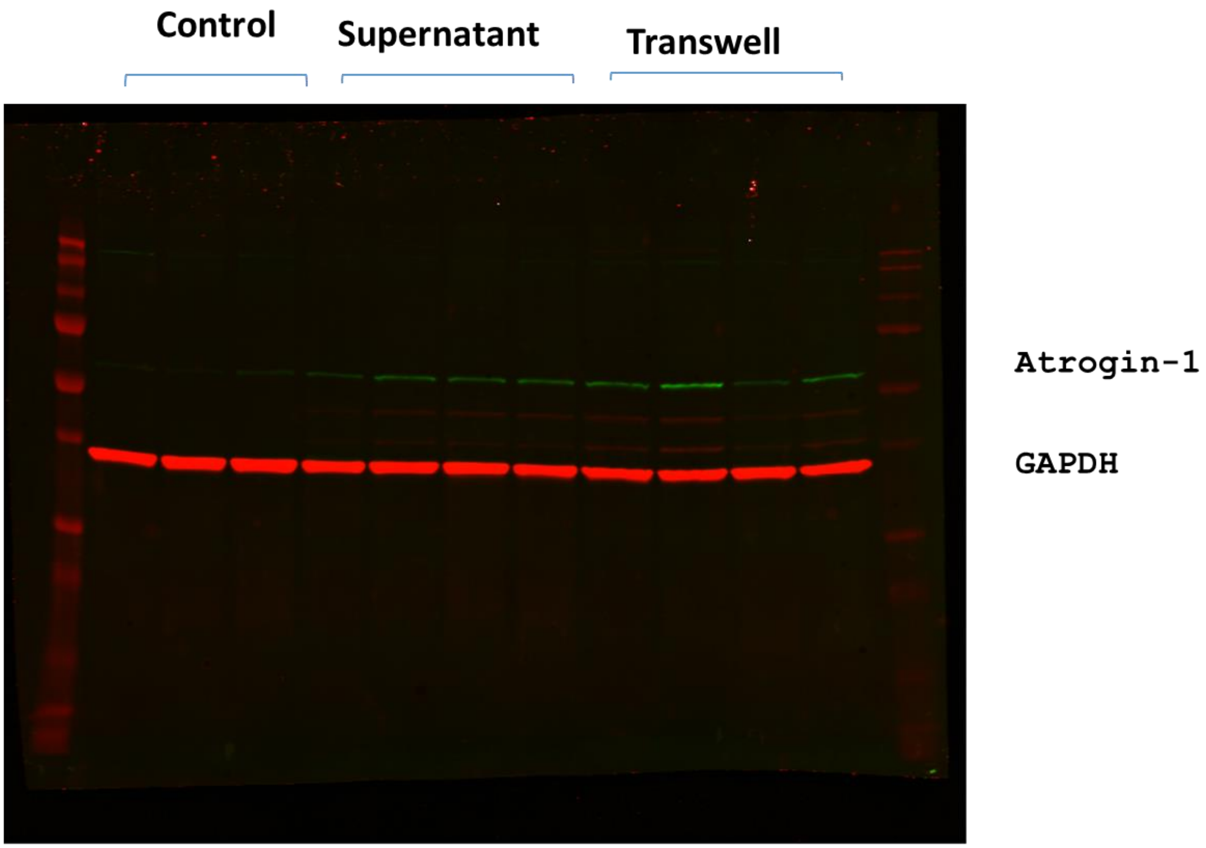

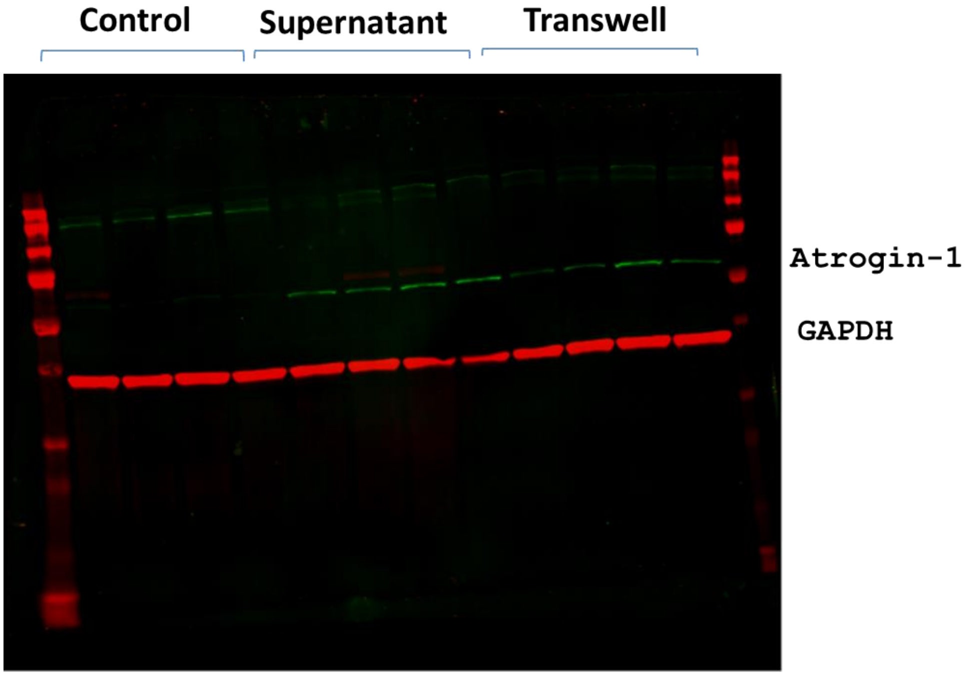
Control
Supernatant
Transwell
Atrogin-1
GAPDH

Whole blots **C26-CM**

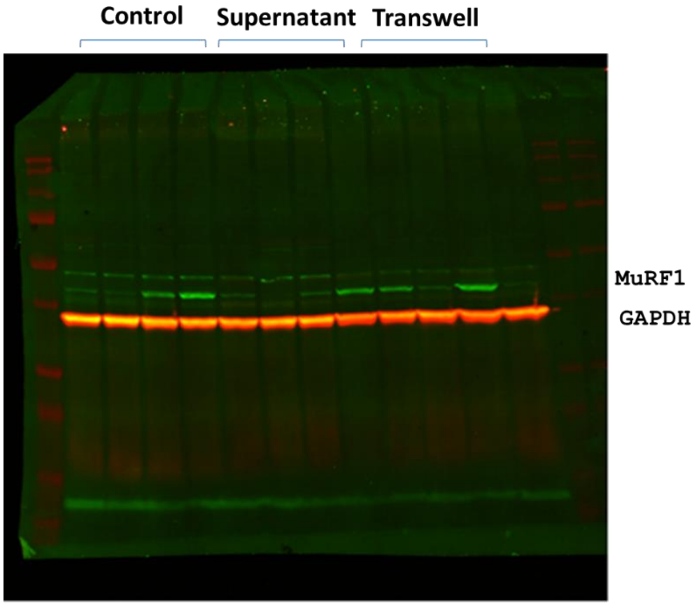


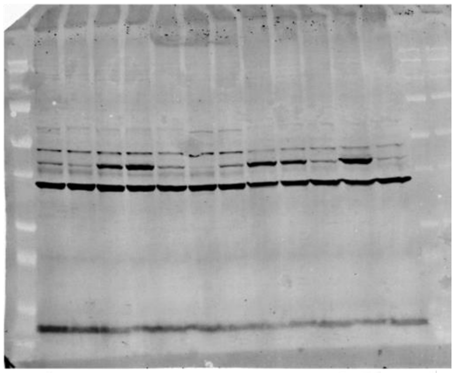

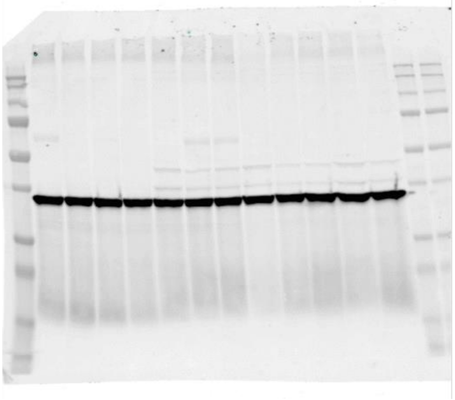

Whole blots – MurF1 + re-stained for LC3B **C26-CM**

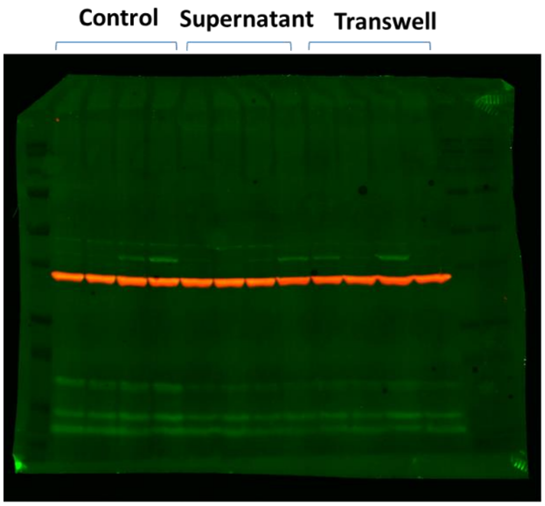


LC3B

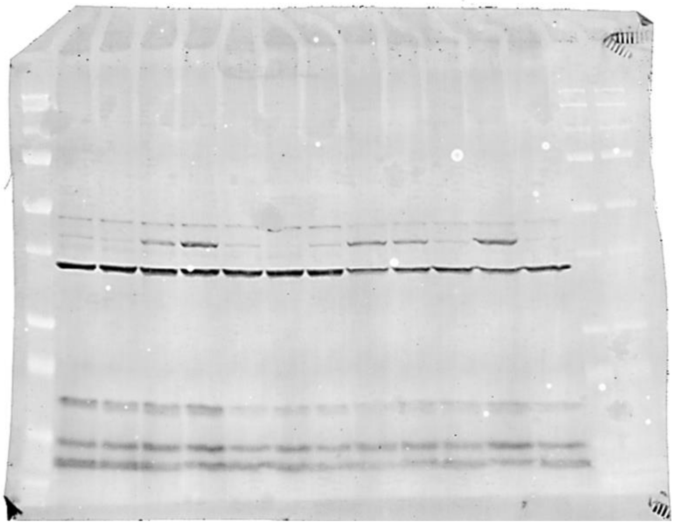

**C26-CM**

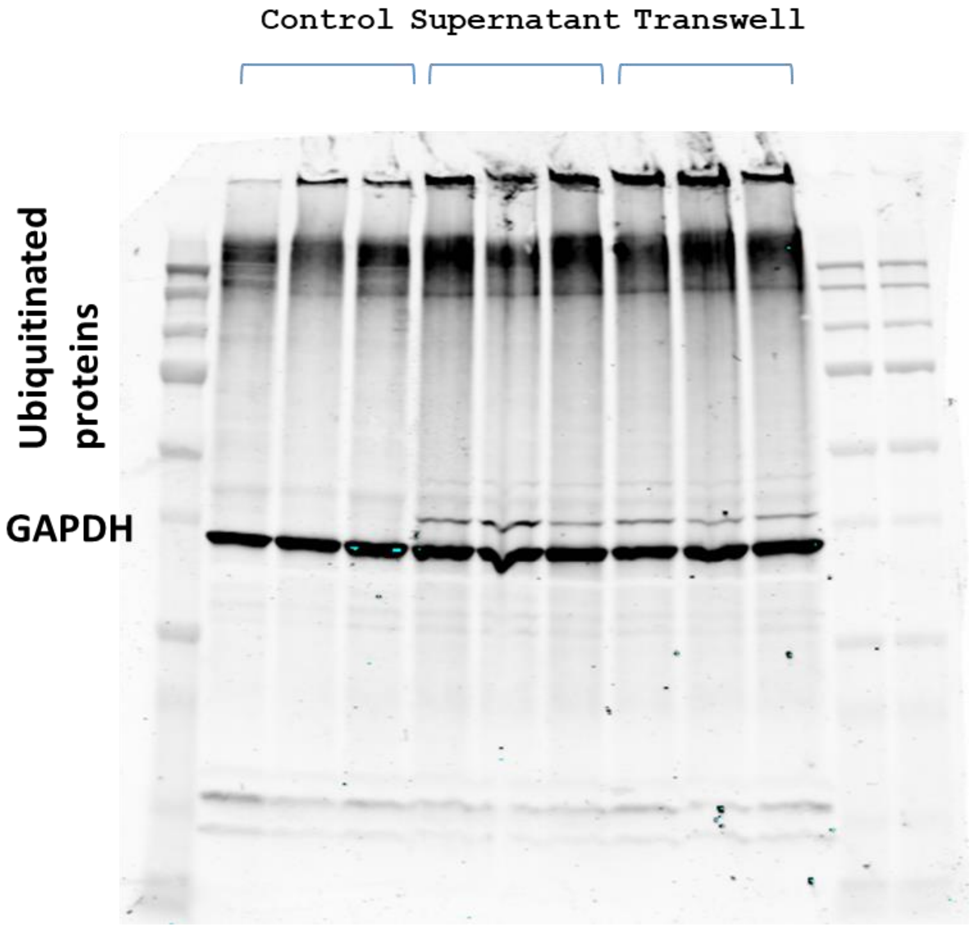
Control Supernatant Transwell
Ubiquitinated proteins
GAPDH

### C26-CM induction – LC3b – Baf A1

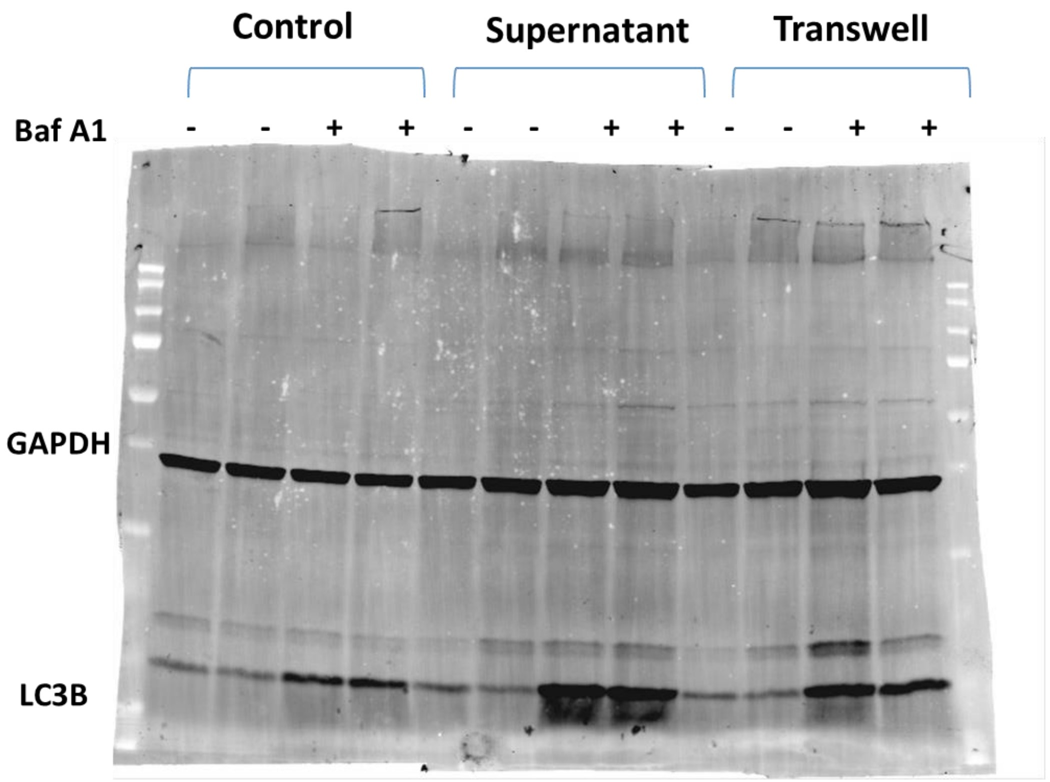

#### C26-CM induction – LC3b – Baf A1

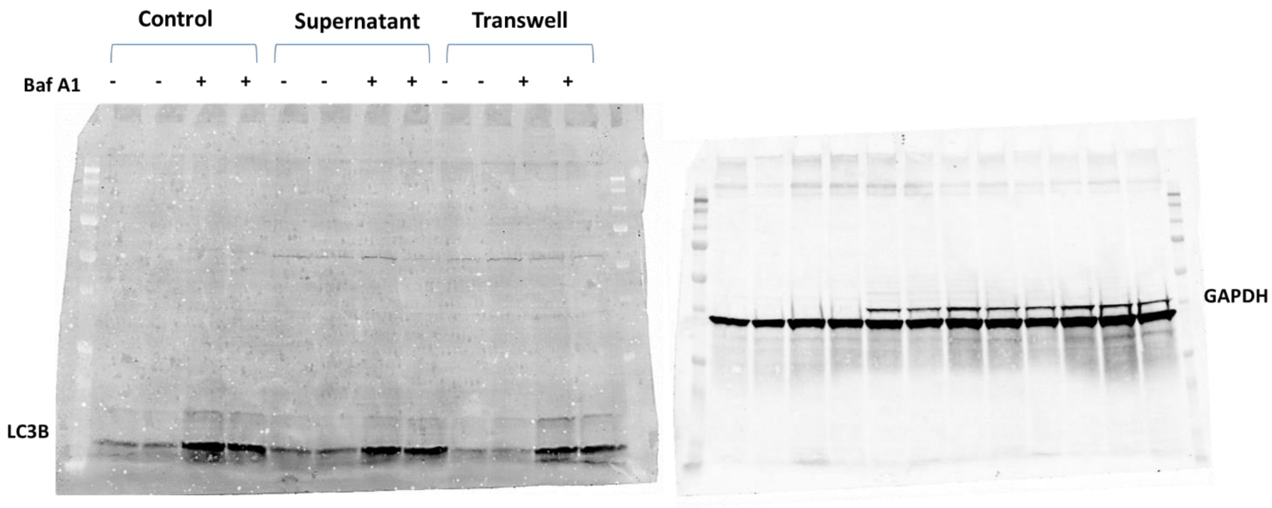

**C26-CM**

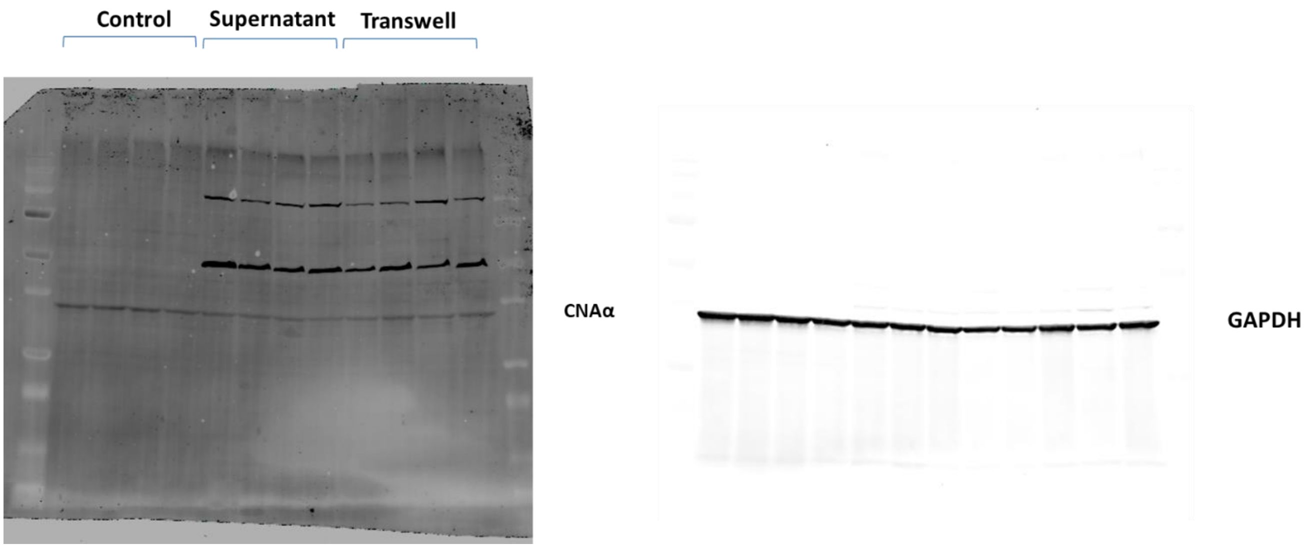

C26-CM

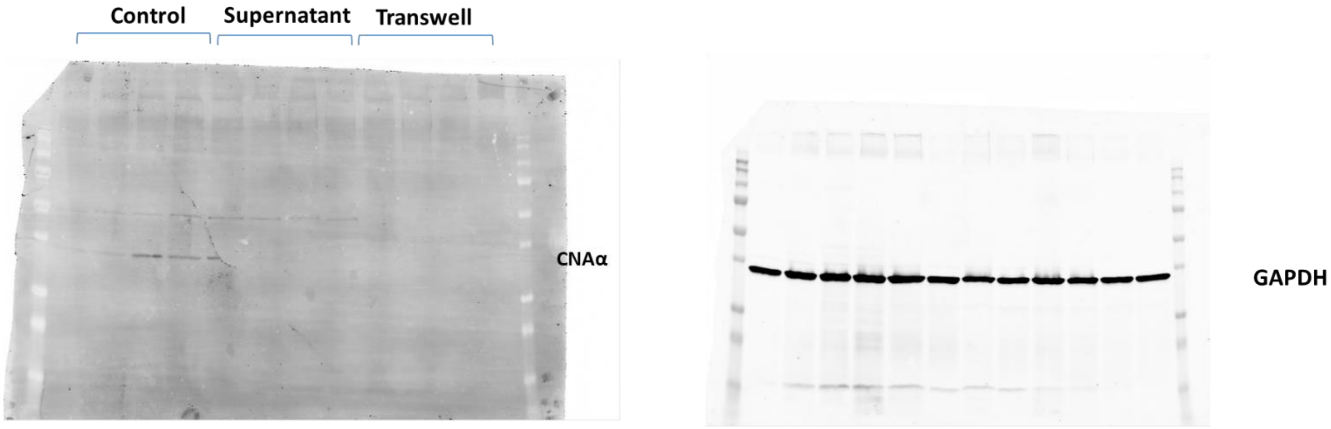

**C26-CM**

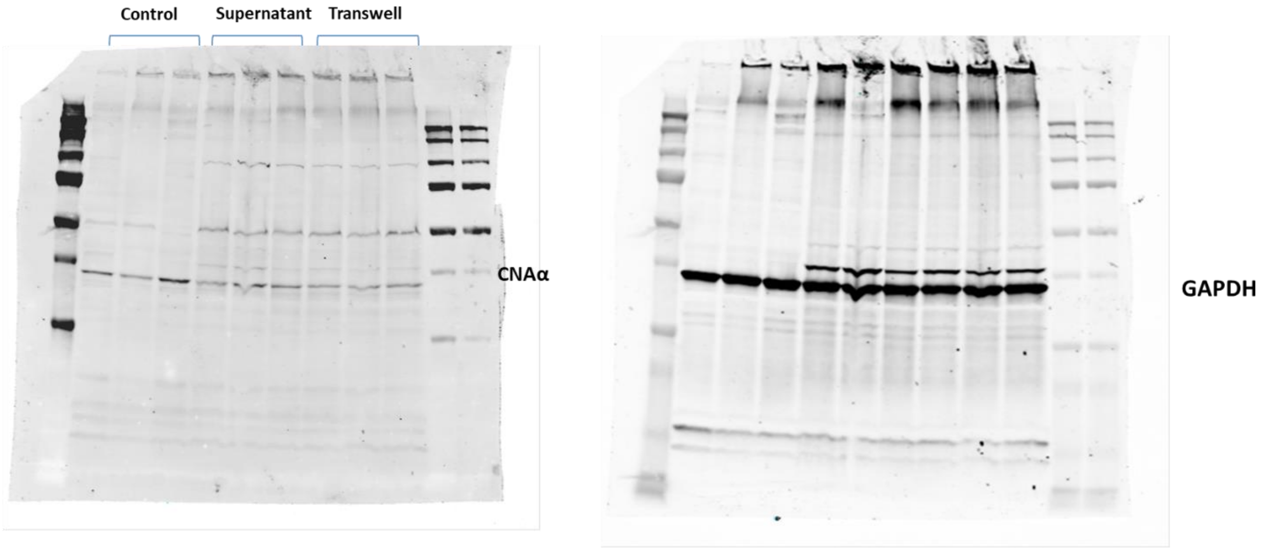

HCT116-CM

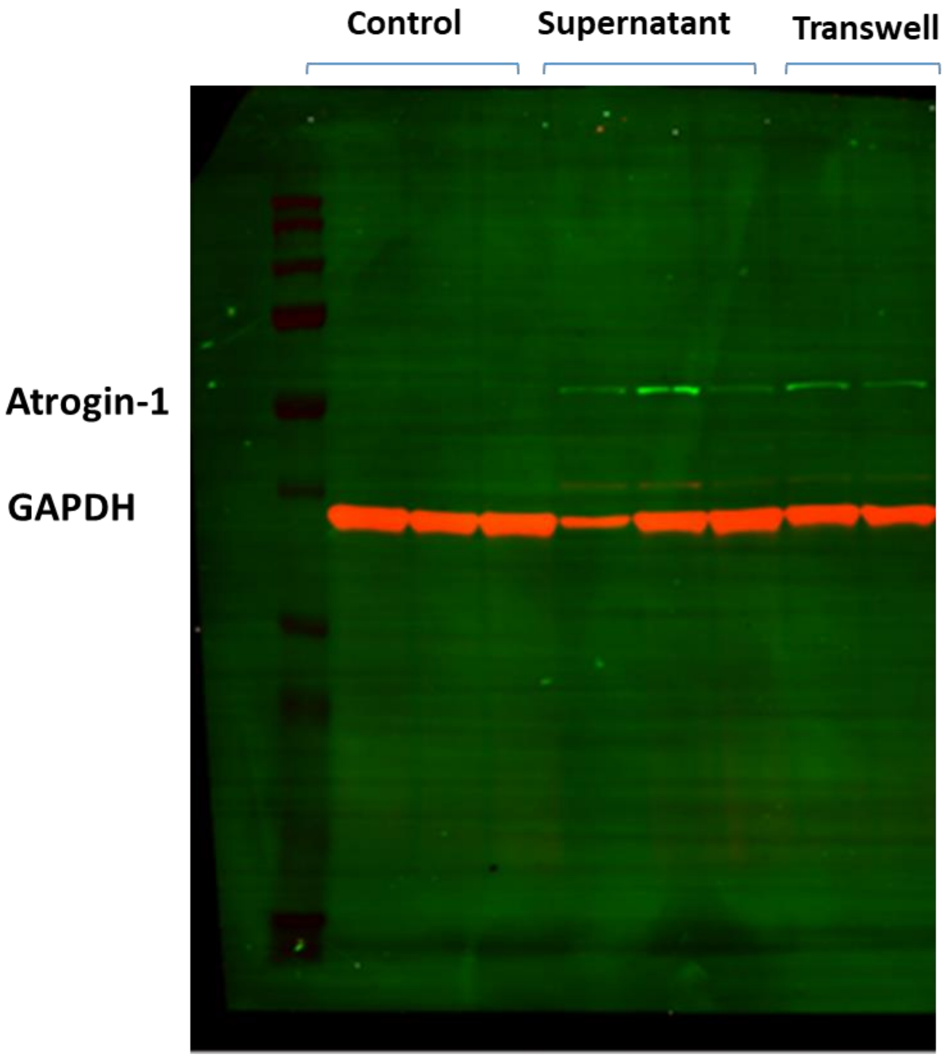

HCT116-CM

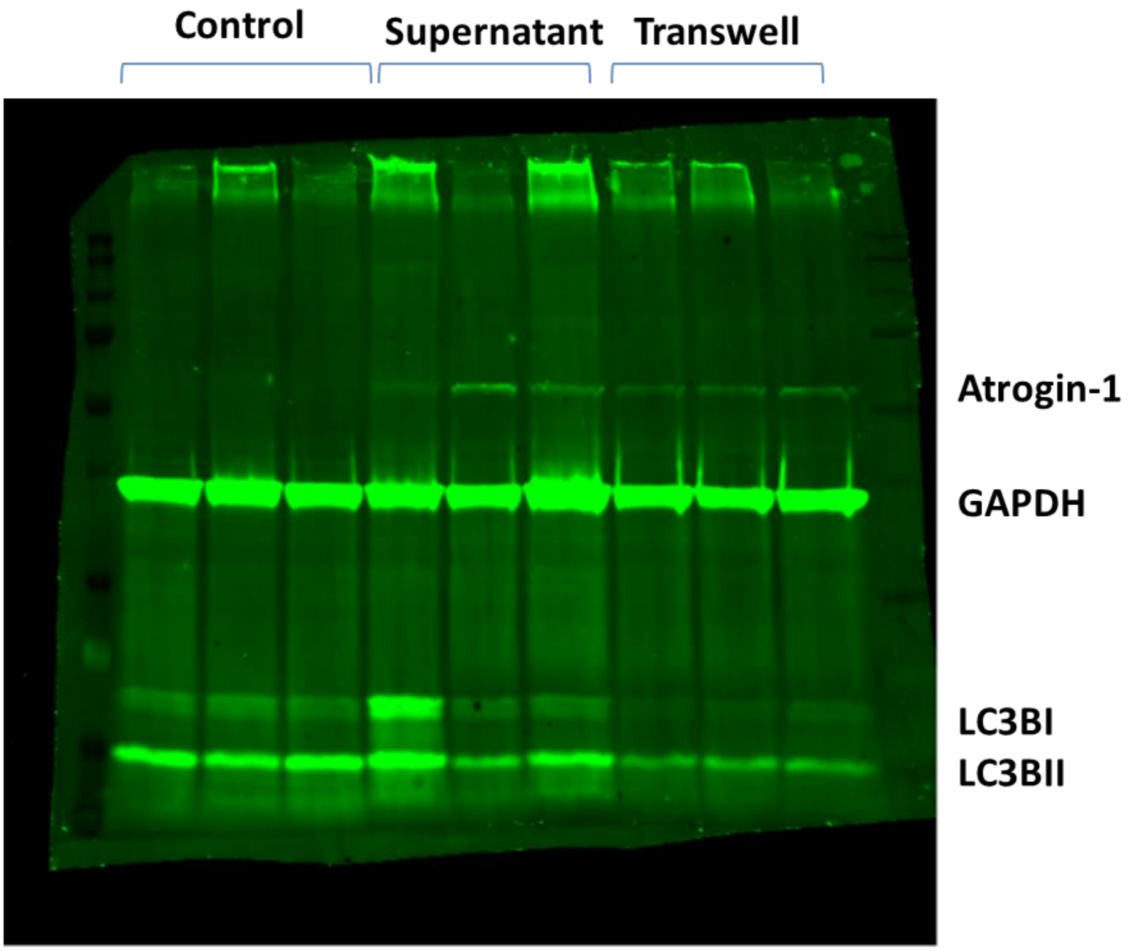
Control
Supernatant
Transwell
Atrogin-1
GAPDH
LC3BI
LC3BII

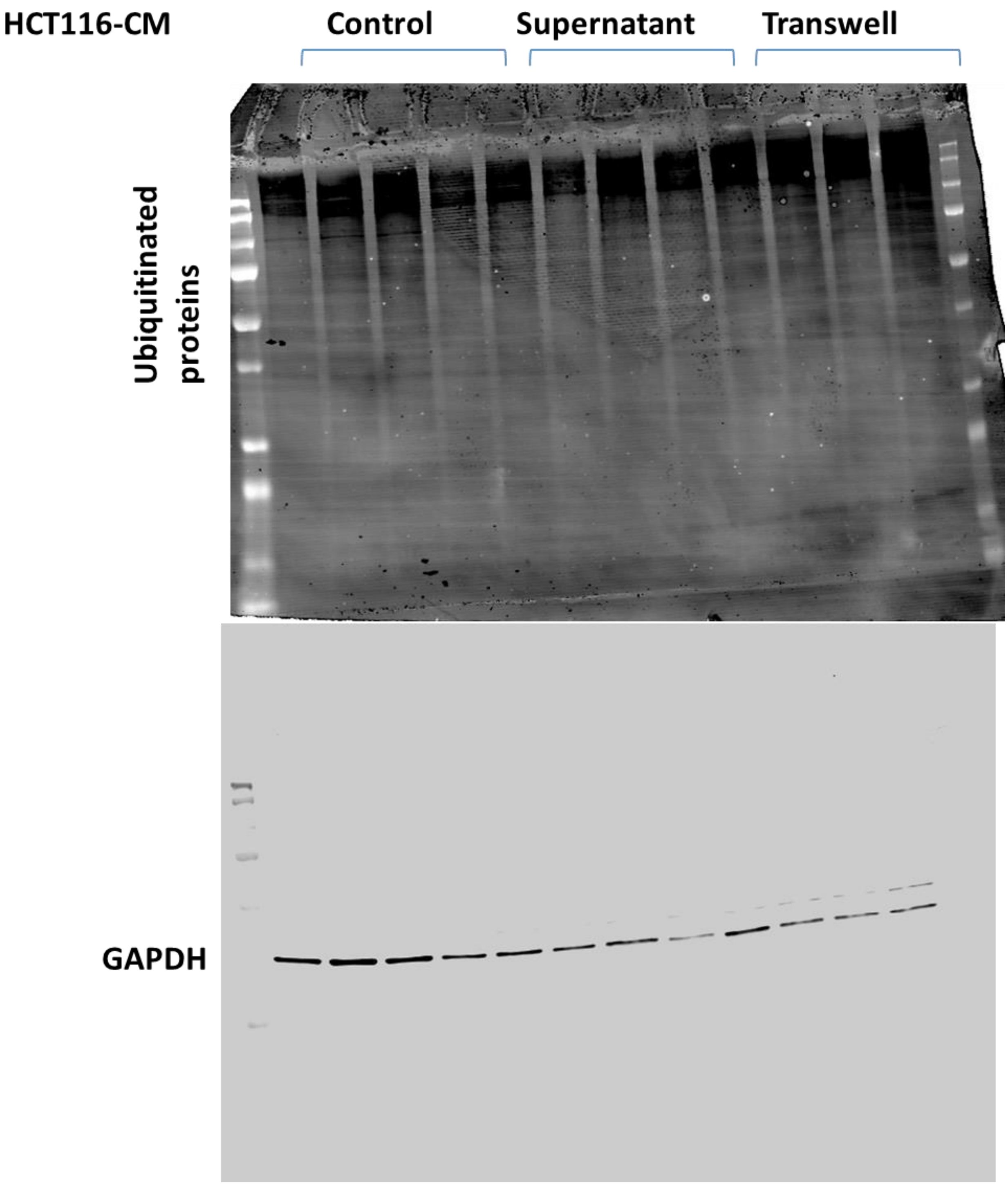
HCT116-CM
Control
Supernatant
Transwell
Ubiquitinated proteins
GAPDH

**HCT116-CM** **α- Calcineurin + LC3B**

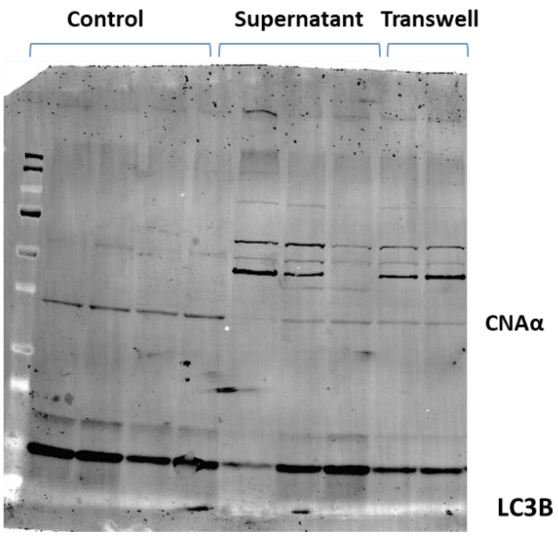


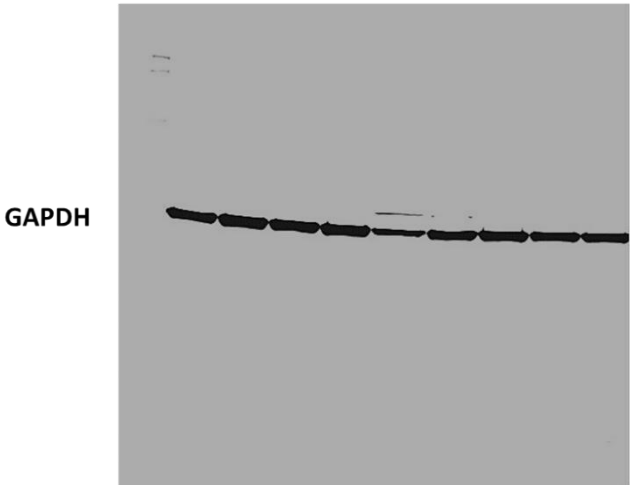

**HCT116-CM**

**α- Calcineurin**

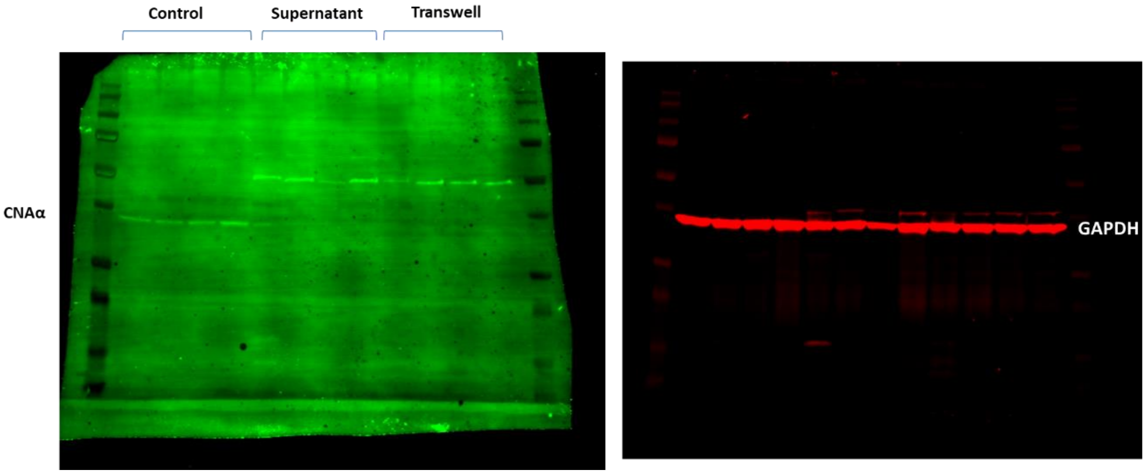

HCT116-CM

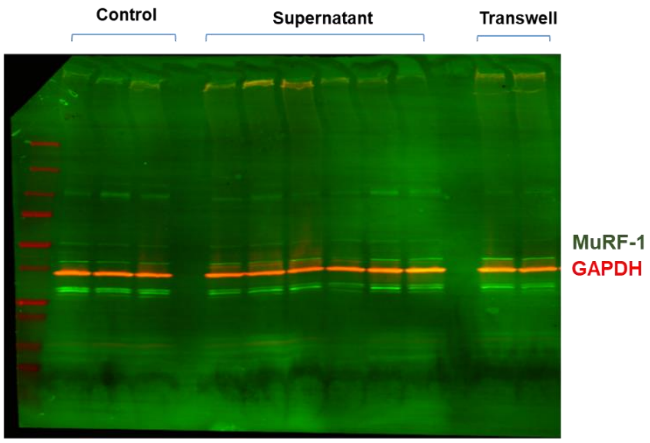


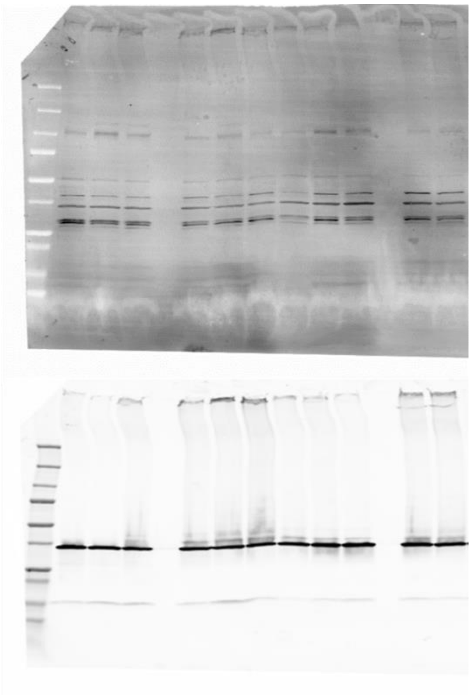

**Cachexia**

**C26 and HCT116-CM**

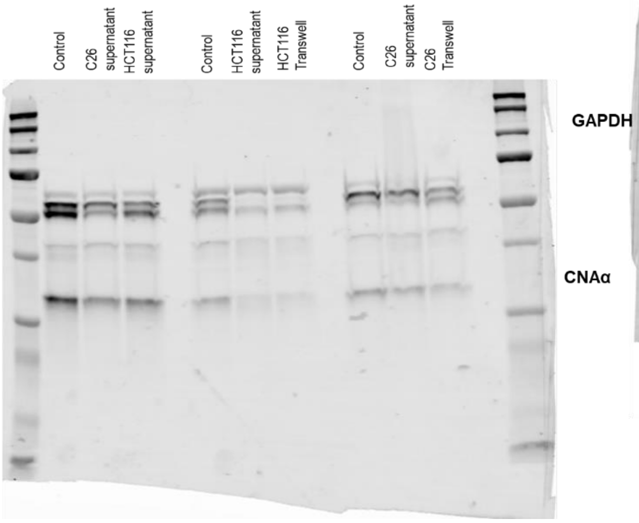


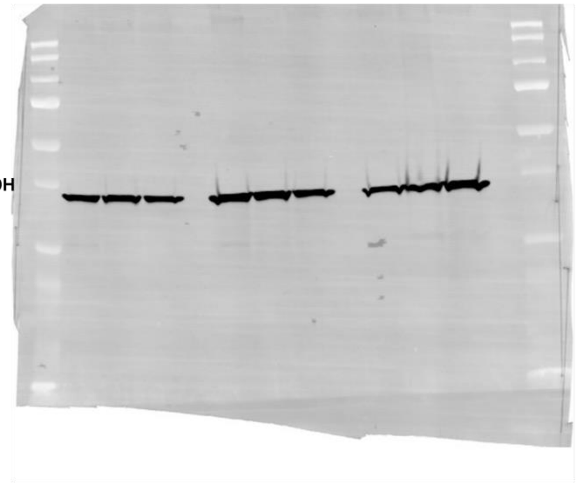

**C26-CM 1week** **Atrogin -1/GAPDH + restained for global ubiquitination**

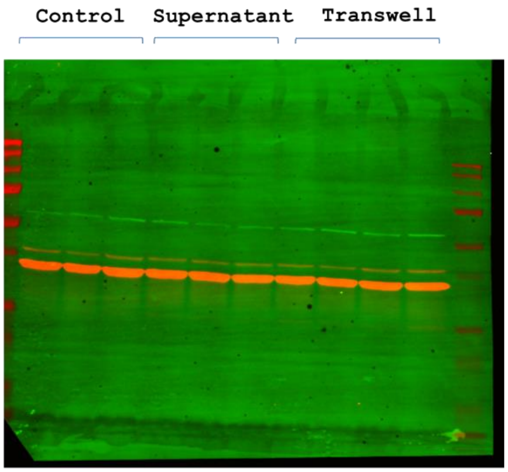


Atrogin-1

GAPDH

Ubiquitinated proteins

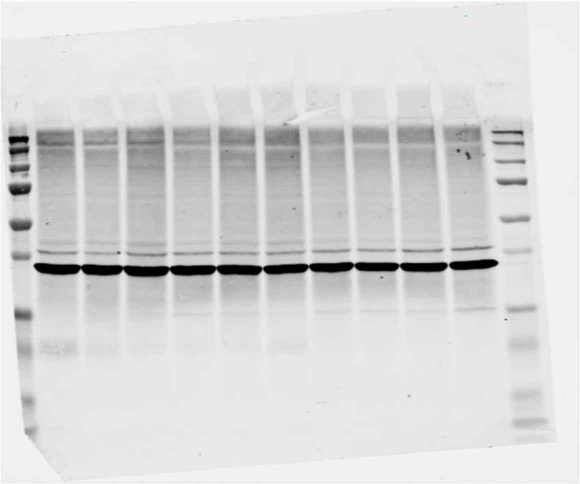

**C26-CM 1week**

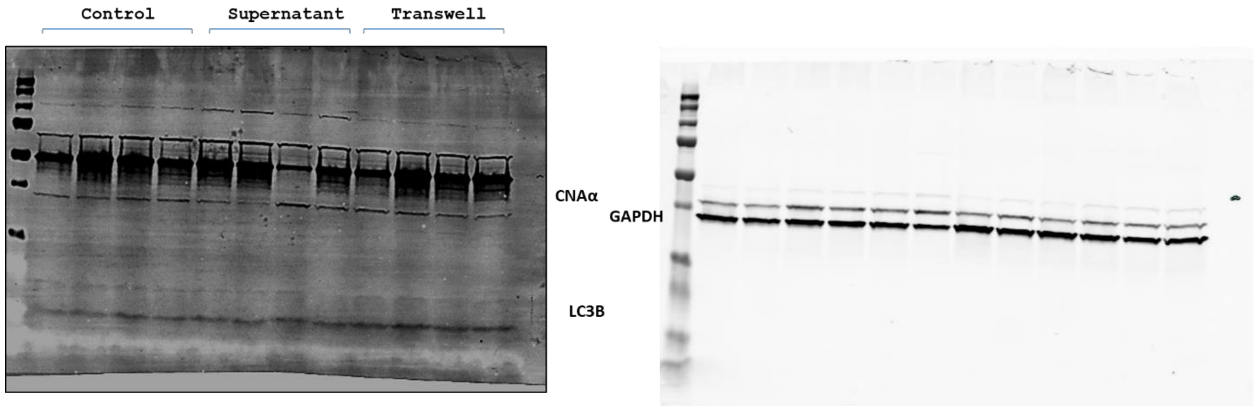

**C26-CM 1week**

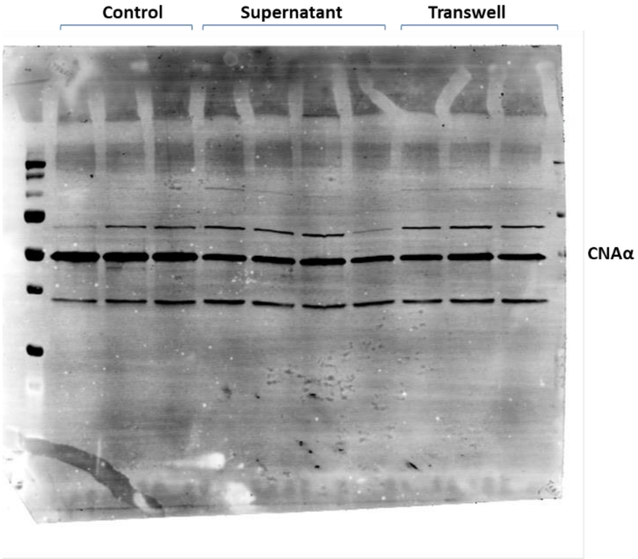


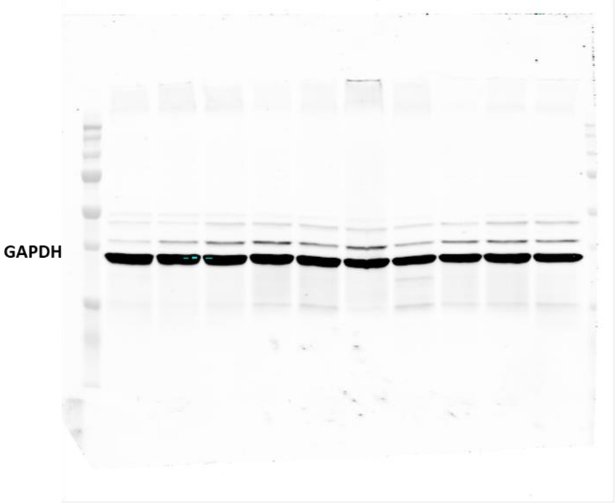

**HC116-CM 1week**

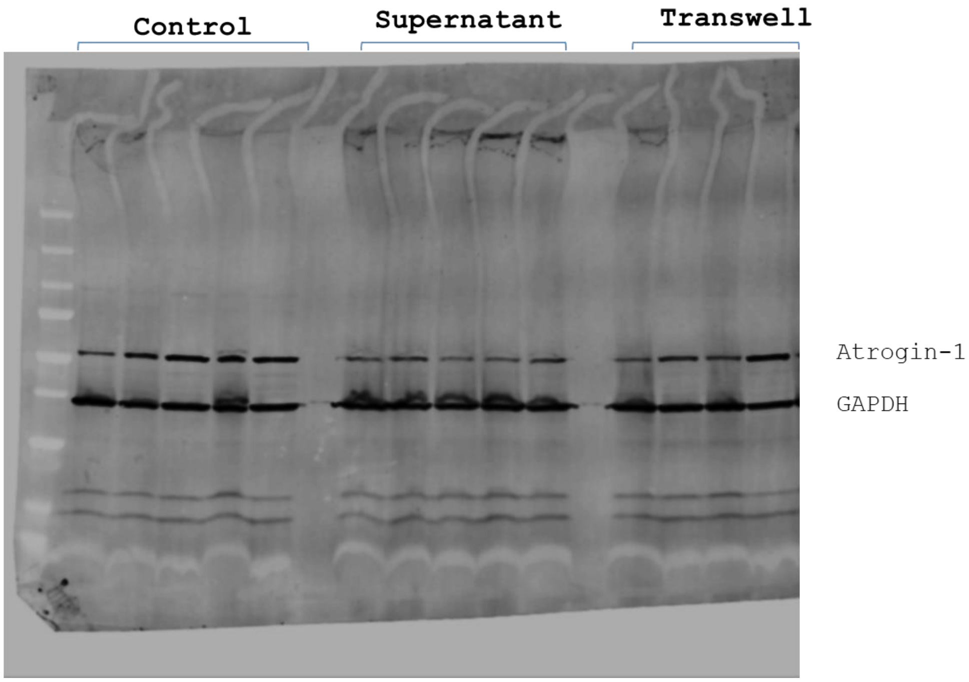

**HC116-CM 1week** **Ubiquitin + restained for LC3B**

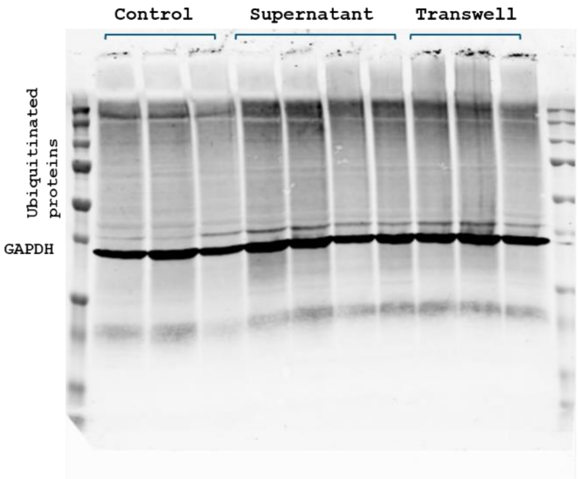


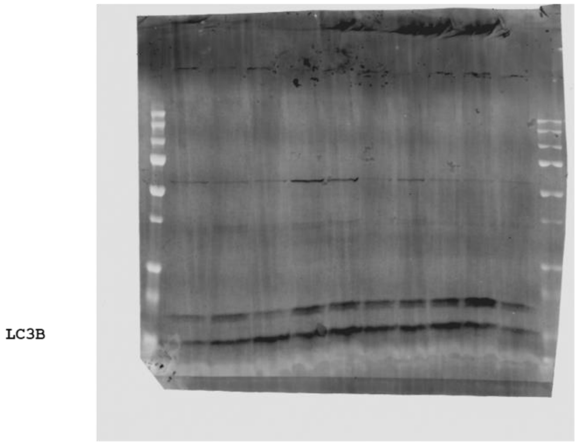

**HC116-CM 1week**

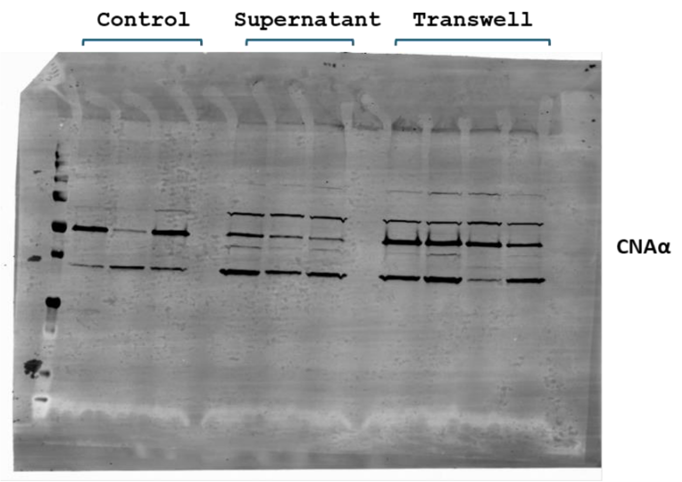


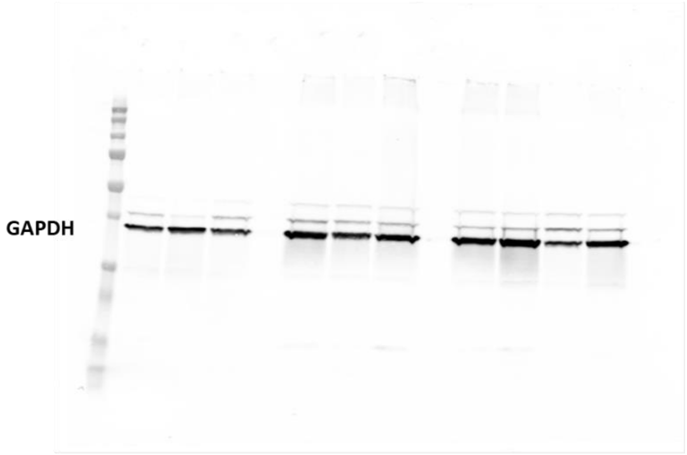


Figure 10.
